# MEG, myself, and I: individual identification from neurophysiological brain activity

**DOI:** 10.1101/2021.02.18.431803

**Authors:** Jason Da Silva Castanheira, Hector D Orozco, Bratislav Misic, Sylvain Baillet

## Abstract

Large, openly available datasets and current analytic tools promise the emergence of population neuroscience. The considerable diversity in personality traits and behaviour between individuals is reflected in the statistical variability of neural data collected in such repositories. This amount of variability challenges the sensitivity and specificity of analysis methods to capture the personal characteristics of a putative neural portrait. Recent studies with functional magnetic resonance imaging (fMRI) have concluded that patterns of resting-state functional connectivity can both successfully identify individuals within a cohort and predict some individual traits, yielding the notion of a *neural fingerprint*. Here, we aimed to clarify the neurophysiological foundations of individual differentiation from features of the rich and complex dynamics of resting-state brain activity using magnetoencephalography (MEG) in 158 participants. Akin to fMRI approaches, neurophysiological functional connectomes enabled the identification of individuals, with identifiability rates similar to fMRI’s. We also show that individual identification was equally successful from simpler measures of the spatial distribution of neurophysiological spectral signal power. Our data further indicate that identifiability can be achieved from brain recordings as short as 30 seconds, and that it is robust over time: individuals remain identifiable from recordings performed weeks after their baseline reference data was collected. Based on these results, we can anticipate a vast range of further research and practical applications of individual differentiation from neural electrophysiology in personalized, clinical, and basic neuroscience.

## Introduction

Understanding the biological nature of individual traits and behaviour is an overarching objective of neuroscience research (*1*–*4*). The increasing availability of large, openly available datasets and advanced computational tools propels the field toward this aim (*5*–*7*). Yet, with bigger and deeper data volumes, neuroscientists are confronted to a paradox: while big-data neuroscience approaches the realm of population neuroscience, we remain challenged by understanding how interindividual data variability echoes the singularity of the self (*1, 3, 8, 9*).

This epistemological question has become particularly vivid with recent research showing that individuals can be identified from a cohort via their respective *neural fingerprints* derived from structural magnetic resonance imaging (MRI) (*10, 11*), functional MRI (fMRI) (*12*–*16*), electroencephalography (EEG) (*17*–*19*), or functional near-infrared spectroscopy (fNIRS) (*20*). Strikingly, neural fingerprints are associated with individual traits such as global intelligence, working memory, and attention abilities (*21*–*24*). Most published work so far is methodologically based on inter-individual similarity measures of functional connectivity—understood as statistical dependencies between ongoing signals across brain regions in task-free awake conditions (*25, 26*)—as defining features of neural fingerprints. Yet, the indirect coupling between hemodynamic and neural brain signaling interrogates the neurophysiological nature of brain fingerprints.

In electrophysiology, ongoing brain dynamics at rest are rich and complex (26) and have long been considered a nuisance, a by-product of neural noise (*28*–*30*). Recent experimental evidence, spurred by systems neuroscience models, indicates that spontaneous brain activity captured using electrophysiological techniques expresses similar resting-state connectomes as fMRI and influences conscious, sensory processes (*31*–*33*). Ongoing neurophysiological activity varies considerably between individuals and across the lifespan. One instance is the inter-individual variability of prominent features of human brain neurophysiological activity, such as the alpha rhythm (8-12 Hz) peak frequency (*34, 35*). Previous EEG fingerprinting work was restricted to scalp data, and therefore, provided limited neuroanatomical insight (*17*–*19*). Another distinctive aspect of electrophysiology is the contamination of recordings by artefacts of different natures including environment and instrument noise, muscle contractions, eye and head movements, which can be distinctive of individuals and can bias fingerprinting with non-neural signal features. Overall, the unique signature components of fast, neurophysiological brain dynamics across individuals remain unchartered.

Here we used resting state recordings of magnetoencephalography (MEG; 27) from a large cohort of participants to identify neurophysiological features of individual differentiation. We derived both measures of functional organization (i.e., functional connectivity) inspired by fMRI *neural fingerprinting* approaches, and spectral signal markers that are proper to the wider frequency spectrum of brain signaling accessible to neurophysiological data.

## Results

We used MEG data from 158 participants available from the Open MEG Archives (OMEGA; 6). Data collected on multiple days were available for a subset of these participants (N=47; mean duration between consecutive sessions: 201.7 days; Figure 1). The participants were both healthy and patient volunteers (ADHD and chronic pain) spanning in age from 18 – 73 years-old (see Supplemental Material). T1-weigthed structural MRI volumes were available from OMEGA for all participants and were used to produce source maps of resting-state brain activity (*36*). We derived several neurophysiological signal features from MEG brain source time series summarized within the Desikan-Killiany atlas—68 regions of interest (ROIs) parcellating the entire cortical surface (*37*). The MEG features comprised power-spectral-density estimates (PSD) within each of the 68 ROIs (*37*), and 68×68 functional connectomes (FC) between these ROIs. The approach is illustrated in Figure 1 and the FC and PSD methodological details are provided in Materials and Methods.

**Figure 1:**
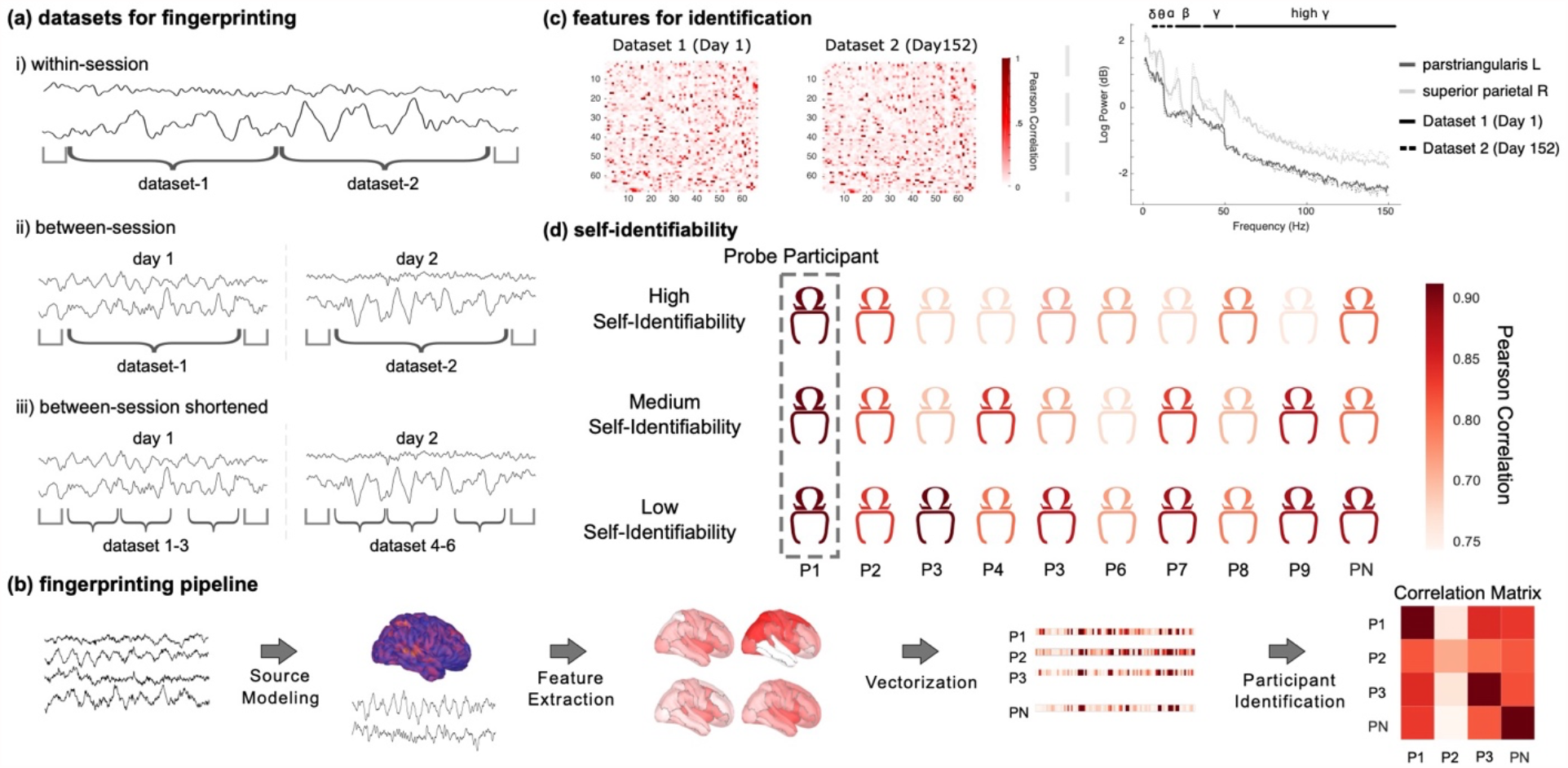
Identification analysis pipeline and definition of self-identifiability **(a)** Schematic of exemplar MEG data divided into datasets used in each of the specified identification challenges. i) *Within-session* challenge: the session data was split in half segments of equal duration; ii) *Between-sessions* challenge: identification was performed using data recorded on two separate days; iii) *Between-session shortened* challenge: data recorded on two different days were split into three 30-s segments. **(b)** Schematic of the data analysis pipeline: source modeling was first performed before extracting features from each region of the Desikan-Killiany atlas (*37*). These features were vectorized and subsequently used to fingerprint individuals, yielding a participant correlation matrix. **(c)** Features for the *between-session* challenge from an exemplar subject. Left panel depicts AEC functional connectivity matrices across two datasets; both matrices feature the Pearson correlation coefficients between all 68 regions of the Desikan-Killiany atlas (*37*). Right panel plots the power spectrum density estimates from two regions of the atlas, across two datasets. **(d)** Self-identifiability was derived for each participant as the z-score of their correlation to themselves, relative to the correlation between themselves and the rest of the cohort. A participant with a high correlation to themselves and low correlations to others was qualified as *highly identifiable*. An individual highly correlated to both themselves and many others in the cohort was qualified as *less identifiable*.

Participant identification was performed across pairs of MEG data segments taken from either the same (*within-session* identification) or a repeated session (*between-session* identification) using two distinct datasets (Figure 1a) and based either on FC or PSD features (referred as *connectome* and *spectral* fingerprinting, respectively). The *within-session* challenge with longer data segments was considered to assess the baseline performances of the MEG fingerprinting approaches proposed. The more challenging situations developed in the present report concern individual identification from shorter 30-s time segments within or between recording sessions. For each pair of participants, the Pearson’s correlation coefficient between their respective features (i.e., FC or PSD) was the corresponding entry in the group correlation matrix (see Supplemental Material). The identification procedure for each individual proceeded via a lookup operation through the corresponding row of the correlation matrix; the index of the column featuring the largest correlation coefficient determined the predicted identity of the individual in the cohort. Thus, if a given individual’s data features from the first dataset were most correlated to the data features from their second dataset, the individual would be correctly identified. Note that taking the maximum along the rows or columns simply switches which dataset is used for deriving the identification features (e.g., identifying individuals using dataset 1 from features derived from dataset 2; results for all possible combinations of datasets are in Supplemental Material). The overall accuracy of the identification procedure was computed as the proportion of participants correctly identified. We ran three types of identification challenges: *within-session* identification consisted of the personal differentiation between 158 participants (i.e., the datasets were from same-day recordings split in half); a *between-session* identification challenge for a subset of 47 participants for whom the datasets were from two separate days; and a *between-session* identification using considerably shortened data segments (30 seconds) (Figure 1a). We conducted the identification challenges using either broadband MEG data or band-limited versions within the typical frequency bands used in neurophysiology. We also derived a self-identifiability score for every participant, which indicates the saliency of the identification of any given individual in the tested cohort (see Material and Methods).

### Within-session connectome and spectral data differentiate individuals

Within-session MEG connectome and spectral fingerprinting achieved 94.9% and 96.2% participant identification accuracy, respectively (Figure 2). This outcome was robust to switching datasets (Supplemental Material). While previous work (*12*) reported that data reduction strategies improved identification performances, this was not the case with our data. Data reduction strategies only marginally improved individual differentiation, as explained in Supplemental Material.

**Figure 2:**
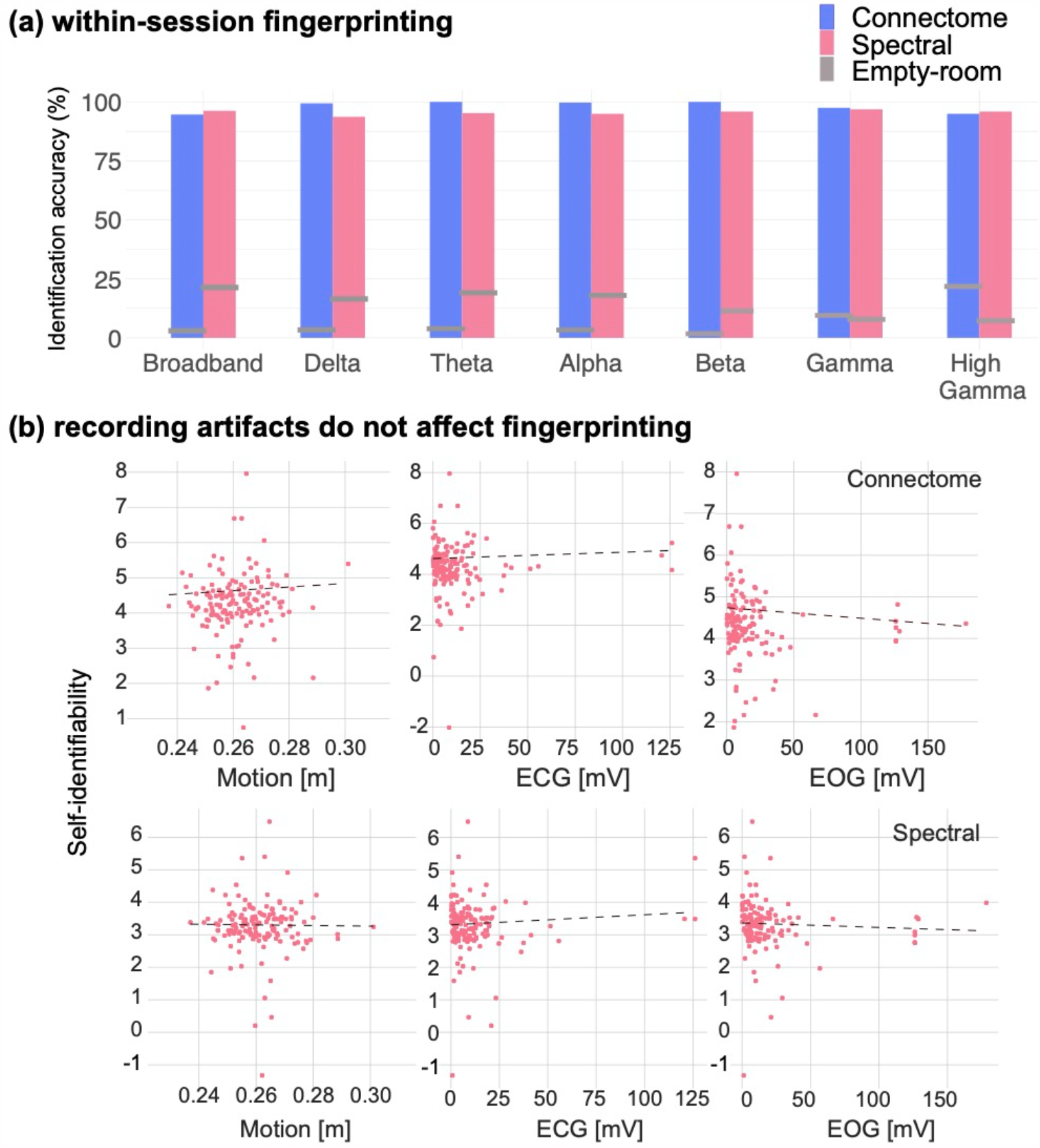
Within-session identification is not related to recording artifacts **(a)** Identification accuracy of connectome and spectral fingerprinting based on broadband and narrowband brain signals. Horizontal grey bars indicate reference identification levels obtained from empty-room data recorded on the same days as participants (see Methods). **(b)** Self-identifiability was not related to typical confounds such head motion, eye movements and heartbeats. Top row: using connectome fingerprinting; bottom row: spectral fingerprinting.

We also ran the identification procedure for each of the typical frequency bands of electrophysiology to understand whether the expression of certain ranges of brain rhythms would be more specific of individual differentiation. We bandpass filtered MEG signals in the delta (1-4Hz), theta (4-8Hz), alpha (8-13Hz), beta (13-30Hz), gamma (30-50Hz) and high gamma (50-150Hz) frequency bands before running the same *within-session* fingerprinting procedure using the resulting narrowband signals. Narrowband connectome fingerprinting yielded identification accuracy scores of 98.7% for delta, 100% for theta, 99.4% for alpha, 100% for beta, 98.7% for gamma, and 94.9% for high gamma. Narrowband spectral fingerprinting produced identification accuracies of 94.9% for delta, 95.6% for theta, 95.6% for alpha, 96.2% for beta, 96.2% for gamma, and 97.5% for high gamma. These results are summarized Figure 2a.

### MEG fingerprinting is robust against physiological, artefactual, and demographics confounds

We investigated the robustness of these results against variables of no interest and possible confounds. We first processed each individual session’s empty-room recordings in an identical fashion to participants brain data. In particular, we produced pseudo brain maps of empty-room sensor data using the same imaging kernels as those used for each session’s participant brain data. The implication is that imaging kernels designed based on information that are specific of each participant, such as their respective head positions in the MEG sensor array and individual anatomy brain features that constrain MEG source maps. We therefore tested whether such individual information unrelated to brain activity contributed substantially to individual identification from MEG source maps. We found that identification performances were considerably reduced using empty-room data (<20% across all tested models; Figure 2). These results based on source maps were corroborated by the low fingerprinting performances obtained by using empty-room sensor data only (<5% across all tested models; Supplemental Material).

We then performed Pearson correlation analyses between identification scores and recording parameters, typical MEG artifacts and demographic variables. There was no association between the duration of scans and self-identifiability for connectome (r=-0.02, p=0.75) and spectral (r=0.02, p=0.8) fingerprinting (Supplemental Material). Further, none of the tested MEG artifacts due to eye movements, heartbeats, and head motion were related to individual identifiability from either connectome or spectral fingerprinting. Indeed, self-identifiability was not correlated to motion (connectome: r=0.06, p=0.5; spectral: r= -0.01, p= 0.9), cardiac (connectome: r=0.05, p=0.6; spectral: r= 0.07, p= 0.4), or ocular (connectome: r= -0.09, p = 0.3; spectral: r=-0.05, p=0.5) artifacts (Figure 2b).

Lastly, we further hypothesized that fingerprinting performances may have been skewed by sample heterogeneity in terms of data from healthy vs. patient participants. Yet, there was less than 1% differences in identification accuracy after restricting fingerprinting to healthy participant data (Supplemental Material). We also verified that participant demographics such as age, sex, and handedness did not contribute to identifiability either (Supplemental Material).

### MEG fingerprinting is robust over time

We tested whether participants who underwent MEG sessions on separate days were identifiable from datasets collected weeks to months apart (with a range of 1 – 1029 days apart and an average of 201.7 days, SD=210.1). We applied the above fingerprinting procedures towards this *between-session* challenge on the subset of participants concerned (N=47). Connectome fingerprinting decreased in performance compared to the identification accuracy scores obtained from the *within-session* challenge (89.4%). Performance of connectome fingerprinting from narrowband signals also decreased, with the greatest robustness obtained from using signals in the beta and theta bands (Figure 3a and Supplemental Material). In contrast, spectral fingerprinting was robust longitudinally, with identification accuracy scores of 97.9% (broadband) and >90% (narrowband) that were similar to those obtained in the *within-session* challenge (Figure 3 and Supplemental Material). Self-identifiability scores were not correlated with the number of days between MEG sessions (connectome: r= 0.09, p = 0.5; spectral: r= 0.08, p=0.65).

**Figure 3:**
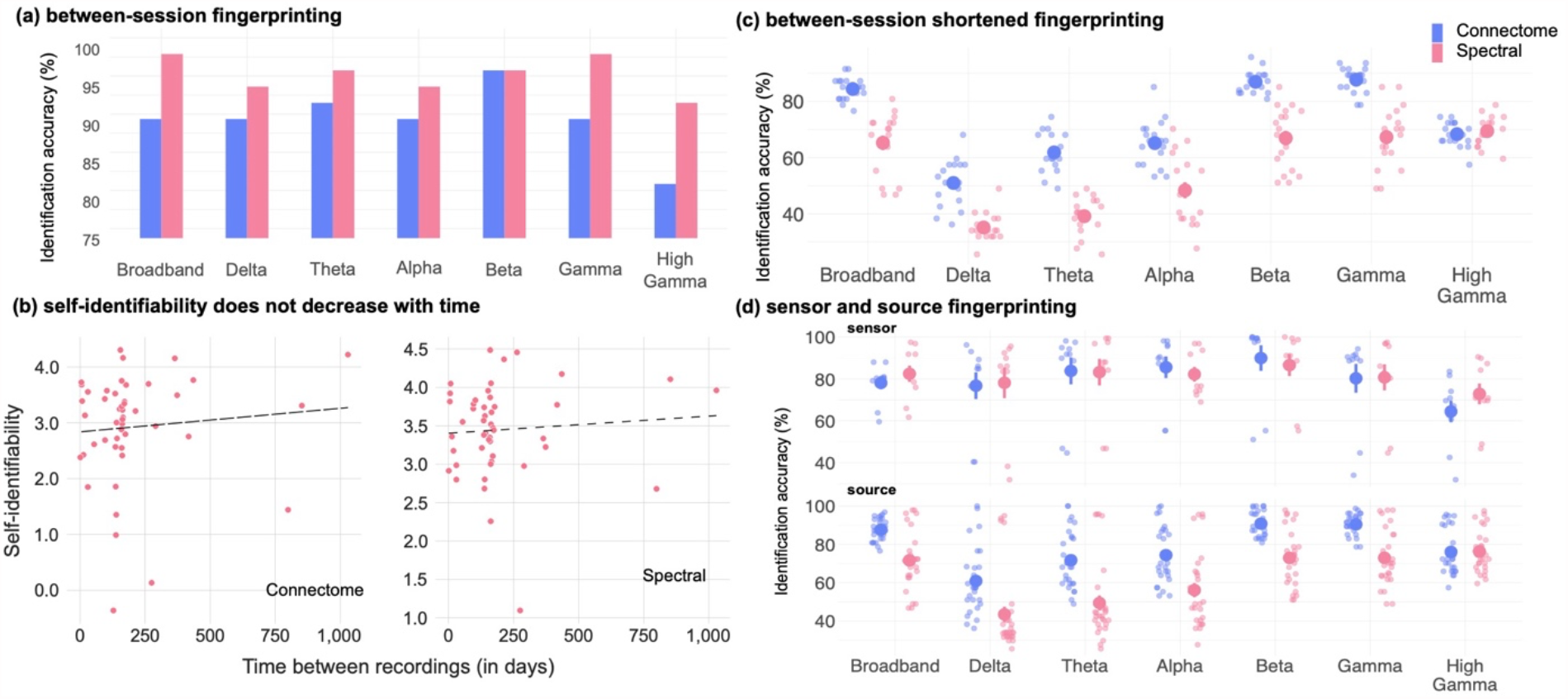
Between-session identification accuracy **(a)** Identification accuracy for connectome and spectral *between-session* fingerprinting. Identification performances are similar to those from the *within-session* challenge. **(b)** Linear regression analyses did not reveal an association between self-identifiability and the delay between session recordings (connectome fingerprinting: r= 0.09, p = 0.5; spectral fingerprinting: r= 0.08, p=0.65). **(c)** *Between-session shortened* identification accuracy using 30-s data segments collected days apart (average: 201.7 days). Each data point represents one combination of datasets used for fingerprinting (see Methods for details) **(d)** Scatter plot of all identification challenges (source and sensor level approaches) across frequency bands for both source and sensor level identification (Supplemental Material details the results obtained for in all sensor data identification challenges.)

We further challenged MEG individual differentiation between sessions days apart using shorter data segments. We extracted three 30-s segments from the *between-session* data on each day (Figure 1a) and ran the same fingerprinting procedures as above. Identification performance from connectome fingerprinting remained high across all 30-s segments tested (Figure 3c) using broadband MEG signals (identification accuracy 84.4%). Performance of spectral fingerprinting was decreased (identification accuracy: 65.2% Figure 3c). We observed similar discrepancies in performance robustness between connectome and spectral fingerprinting using narrowband signals (Figure 3), especially in the delta, theta, and alpha bands. We report results obtained from using sensor data only and for the *within-session shortened* challenge in Supplemental Material.

### Salient neurophysiological features for identification

We identified the features which were the most characteristic of individuals for MEG fingerprinting. We derived measures of intraclass correlation (ICC) (*12*) to quantify how much each feature, such as an edge of the FC connectome or the signal power in a frequency band from an anatomical parcel, contributed to fingerprinting (see Methods). This metric was reported in previous brain fingerprinting studies and captures the inter-rater reliability of each participant as their own rater, to identify the neurophysiological signal features that are the most consistent across individuals (*12, 38*). We performed this analysis for both the broadband connectome and the band-specific spectral fingerprinting *within-session* challenges. The data show that the dorsal attention and visual networks were the most specific across individuals for connectome fingerprinting, in all frequency bands (Figure 4). Beta-band connectivity of the limbic network was particularly distinctive of individuals. For spectral fingerprinting, beta, gamma, and high-gamma band signal power were the most salient identification features, especially across medial regions (Figure 4b). Particularly, signals in the theta, alpha, beta, and gamma bands discriminated individuals along midline, parietal, lateral temporal, and visual areas. These results are consistent with our narrowband analysis (see Figure 2a), which highlights beta activity as the most informative in identifying individuals.

**Figure 4:**
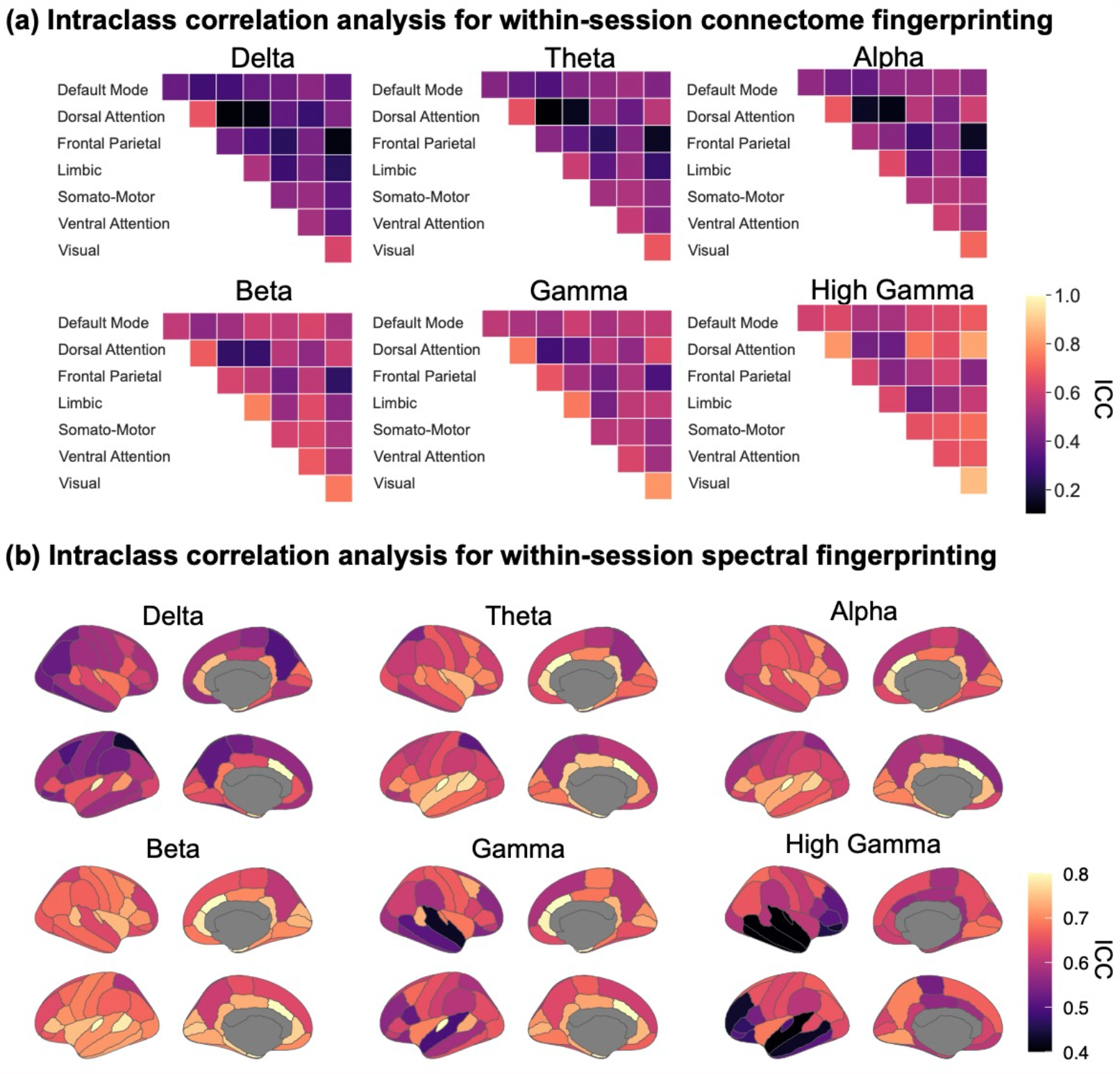
Characteristic features of connectome and spectral fingerprinting Intraclass correlation (ICC) for connectome and spectral *within-session* fingerprinting. **(a)** ICC for connectome fingerprinting plotted for each tested frequency band, using network labels from Yeo et al. (2011). The most prominent networks for connectome fingerprinting were the Visual, Dorsal Attention and Limbic networks. **(b)** ICC for spectral fingerprinting plotted for each tested frequency band and mapped using the Desikan-Killiany cortical parcellation (*37*). The most salient features were the gamma and high-gamma band signals expressed in midline structures and the beta band across the cortex.

### Neurophysiological identifying features are associated with demographics

Beyond identifying individuals in a cohort, we tested whether resting-state neurophysiological features could also predict meaningful participant traits, using an exploratory partial-least-squares (PLS) analysis (see Methods; (*39*)). Briefly, PLS explains the structure of the covariance between two observation matrices – here a demographic matrix and a neurophysiological signal matrix composed of ROI-specific connectome of spectral measures – with latent components. PLS analysis of our data revealed three significant latent components, which were distinct for connectome and spectral fingerprinting (Supplemental Material). The first latent component in connectome fingerprinting was related to clinical population (r= 0.2, 95% CI [0.160, 0.3]) and handedness (r= 0.2, 95% CI [0.1, 0.3]). This demographic profile was associated with reduced beta-band functional connectivity over the frontal parietal network (Figure 5). For spectral fingerprinting, the first salient latent component was related to a younger age (r= -0.3, 95% CI [-0.1, -0.5]), female (r= 0.4, 95% CI [0.2, 0.5]) and clinical population (r= 0.5, 95% CI [0.2, 0.5]). This demographic profile was associated with stronger expressions of broadband neurophysiological signal power in superior parietal regions and the pericalcarine gyrus bilaterally, and reduced neurophysiological signals in the isthmus cingulate (Figure 5).

**Figure 5:**
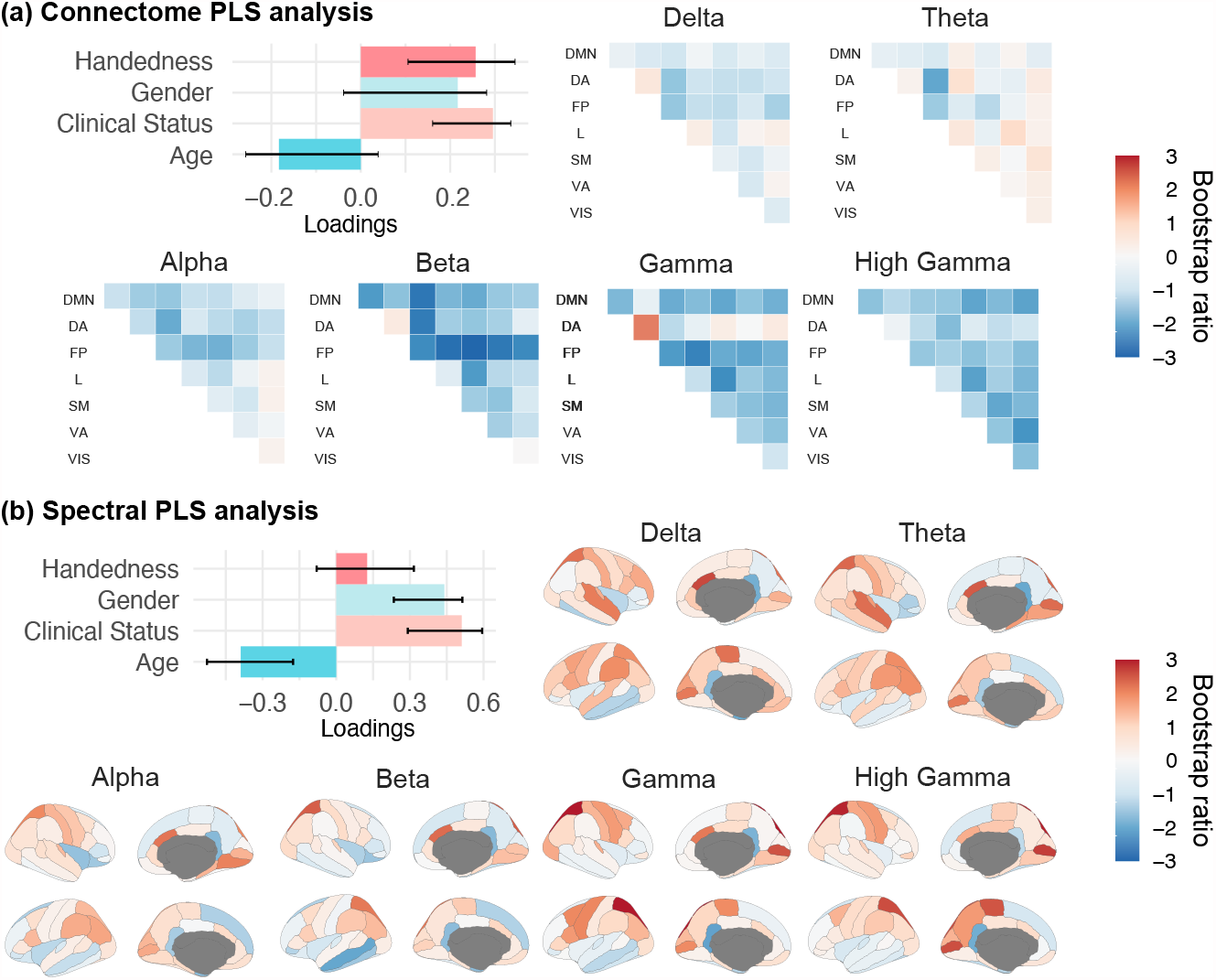
Partial Least-Squares analysis relates demographics to connectome and spectral features **(a)** and **(b)** from left to right, depicts the design saliency patterns for the first latent variables and their associated neural-data bootstrap ratios. Confidence Intervals (CI) were calculated through a bootstrapping procedure, and as such may not necessarily be symmetric. Bootstrap ratios computed for **(a)** connectome and **(b)** spectral features are plotted according to the resting-state networks labelled according to Yeo et al. (2011) and the Desikan-Killiany parcellation (*37*), respectively: Default Mode Network (DMN), Dorsal Attention (DA), Frontal-Parietal (FP), Limbic (L), Somato-Motor (SM), Ventral Attention (VA), and Visual (VIS).

## Discussion

The recent leveraging of large, open fMRI datasets has brought empirical evidence that individuals may be identified within a cohort from their brain imaging functional connectome, inspiring the metaphor of a *neural fingerprint*. Unlike hand fingerprints, their cerebral counterpart predicts task performance and a variety of traits (*14, 21*–*24*). These intriguing findings require a better understanding of their neurophysiological foundations, which we sought to characterize from direct neural signals captured at a large scale with MEG. Our data show that individuals can be identified in a cohort of 158 unrelated participants from their respective resting-state connectomes and spectral profiles in a range of fast brain signals. MEG fingerprinting was successful using data lengths (30 seconds) much shorter than those reported for fMRI fingerprinting (*14, 41*). Brain electrophysiological signals are rich, complex and convey expressions of large-scale neural dynamics channeled by individual structural anatomy and physiology (*42*). Indeed, we also showed that MEG fingerprinting is robust across time, making individuals potentially identifiable from data collected days, months, or years apart.

Lastly, we characterized whether individual differences in resting-state neural dynamics are demographically meaningful through an exploratory PLS analysis. We showed that both resting-state functional connectomes and spectra predict latent demographic components. Recent findings corroborate our results, demonstrating individual differences between functional connectomes derived from resting-state electrophysiology (*43*). Future work will be required to replicate and expand these findings in more samples of individuals.

### Connectome and spectral neurophysiological fingerprints

Our results highlight two sets of brain-wide electrophysiological features that contributed to successful individual identification: connectome and spectral measures across the neurophysiological frequency spectrum. Overall, connectome and spectral fingerprinting with MEG performed equivalently to fMRI approaches, achieving overall identification rates above 90%, with robust individual identification over time and against noise (*12, 14, 44*). We found that for connectome fingerprinting, the anatomical regions the most characteristic of individuals differed between MEG and fMRI. While fMRI highlighted the default-mode network and the fronto-parietal resting state networks, MEG connectome fingerprinting emphasized functional connectivity within limbic and visual networks as contributing to individual specific neurophysiological signatures. In contrast, both MEG and fMRI fingerprinting emphasize the importance of the dorsal attention network (*14*). These observations are not mutually exclusive, considering the different nature of brain signals captured by the respective modalities. One possible interpretation—requiring further investigation— is that the fast neurophysiological signals that contribute to identification with MEG have hemodynamic counterparts that are not as salient in fMRI as the identifying networks reported so far. Nevertheless, our data indicate that neurophysiological signals in the beta band contribute to the highest identification accuracy amongst all other typical bands. This finding is compatible with previous work reporting that correlated amplitude changes of MEG brain signals are related to the microstructure of white matter tracts and reveal, with the same amplitude envelope correlation method as used here, MEG resting-state brain networks that align with fMRI’s (*45, 46*). Beta-band activity also emerges from recent literature as a signalling vehicle of re-afferent “top-down” communications in brain circuits (*47, 48*). One can therefore speculate that beta-band signals would convey electrophysiological representations of internal cognitive models that are by essence intimately specific of each individual (*27*).

Such brain signal amplitude signatures are further emphasized by the ability of simple spectral brain maps to enable MEG fingerprinting. *Within-* and *between-session* spectral identification were achieved with remarkable accuracy (>90%) with broadband MEG brain signals or restricted to the typical bands of electrophysiology. Spectral identification based on signals from the faster bands (gamma and high-gamma) was overall the most robust longitudinally and against using shorter data segments. This observation is consistent with the width of (high) gamma frequency bands spanning broader ranges (here between 30-50 Hz and 50-150 Hz) than slower bands such as delta (1-4 Hz), theta (4-8 Hz) and alpha (8-12 Hz). The spectral estimates averaged across the broader (high) gamma bands were therefore the most robust against using shorter data segments. The reduced number of sliding time windows available over shorter data durations increased the variance of the summary statistics extracted to derive the spectral fingerprints from the signals defined over narrower bands. The higher frequency bands were less affected because the larger number of frequency bins involved in the extraction of their summary power statistics tended to compensate the higher empirical variance of spectral estimates from a lesser number of observations over time. Connectome fingerprinting was more immune against using shorter data durations. The underlying approach indeed did not require spectral transformations but resorted to a bank of narrowband filters applied over the original duration of MEG recordings, before the resulting filtered signals were segmented in shorter epochs for the identification challenges. The consequence is that the number of data points used for all narrowband signals was identical across all frequency bands, yielding moderate variability in identification performances compared to those obtained with the spectral approach. Another factor of robustness of the connectome approach is that connectivity weights between network nodes may fluctuate very slowly over time in task free brain activity: Florin and Baillet (31) reported fluctuation rates of 0.01Hz in MEG, indicating typical time cycles of 100s — a duration substantially longer than the 30-s shortest time window used here. Over longer periods of time though, such as in the *between-session* challenge, spectral fingerprinting outperformed its connectome counterpart. We note a slight increase of spectral identification accuracy in the *between-session* challenge (e.g., +1.6% for broadband fingerprinting) compared to *within-session*, which was a statistical fluctuation due to using a smaller sample of participants.

On average across all source fingerprinting challenges reported herein, and despite successful identification across lower frequency bands (delta 52.2%, theta 60.6%, alpha 65.3%), performances were markedly better using high-frequency signal components (beta 81.9%; gamma 81.7%; high gamma 76.2%). Gamma and faster activity have long been associated with concurrent and colocalized hemodynamic fluctuations (*49, 50*). Because they may be seen as dual manifestations of BOLD signaling used in fMRI fingerprinting, this may explain why these signals contributed robustly to MEG brain fingerprinting in our data. However, gamma-band and faster brain signals are on average weaker in amplitude and therefore may be masked by contamination from artifacts and noise (*51*–*53*). The preprocessing applied to our data attenuated such nuisance to a point where individuals were not identifiable from typical sources of signal contamination such as individual head motion behavior.

Although a rhythm of prominent amplitude in humans during rest, alpha-band activity (8-12Hz) was not particularly specific to identify individuals in the cohort. In that respect, our data is aligned with previous MEG works on resting-state connectomes extracted from neurophysiological MEG signals, which did not report on a salient role of alpha activity in driving inter-regional connectivity (*31, 45*). We argue that the spatial topography of alpha resting activity may be relatively stereotypical across individuals, involving thalamo-cortical loops that project focally to the parieto-occipital junction, with limited variability across individuals (*6*). In task, alpha activity has been related to attention orienting, alertness and anticipation, and the registration of (multimodal) sensory information, thereby reflecting transient mental states (*41, 54*–*57*) rather than individual traits.

The data also indicates that MEG fingerprinting is robust against typical recording artefacts that may be idiosyncratic of individuals and therefore, could have confounded identification. In particular, session environmental conditions captured by empty-room MEG recordings were not sufficient to identify individuals within or between sessions. The participant’s anatomical and head-position information embedded in their respective MEG source imaging kernels were also not sufficient to identify individuals. Note that head position changed between sessions. Further studies are required to clarify how these results may vary depending on the type of MEG source modelling adopted. We anticipate little influence of the type of source model used though, based on evidence that beamforming kernels are mathematically equivalently to other major classes of linear source estimation kernels, such as weighted minimum-norm estimators (*58*). Future work should corroborate these results with regards to fingerprinting. The choice of connectivity measure to derive electrophysiological connectomes may also influence identifiability (*59*). We look forward to current progress in electrophysiological brain connectomics to put forward measures of network connectivity informed by mechanistic principles and emerging as a standard metrics in the field to confirm and expand present fingerprinting results (*60*).

While our present data show robust longitudinal fingerprinting performances, future work involving more participants with multiple MEG visits is required to both replicate these observations and investigate whether individual deviations from baseline fingerprints could be early signals of asymptomatic neuropathophysiology (*27*). We hope the remarkable ability to fingerprint individuals from the present electrophysiological features serves as a steppingstone for future investigation, which may include multimodal non-invasive assessments based on MEG, possibly combined with e.g., fMRI and/or EEG.

### Neural fingerprints of individual traits

Our data suggests that individual differences in resting-state neurophysiological functional connectivity and spectral power relate to latent demographic clusters. These observations are in line with previous fMRI work that showed that connectomes are predictive of individual differences in attention, working memory and intelligence. For instance, connectivity patterns between the default mode and the dorsal attention networks predict attentional behaviour during task and self-reported mind wandering (*22, 61*, see *62 for review*). Overall, a possible conceptual framework is that task free neural dynamics are the signatures of an individual scaffold of brain functions that is predictive of task behaviour. This view is also that of the spontaneous trait reactivation hypothesis wherein the organization of the human cortex at rest (manifested e.g., by functional connectivity) is a window into the self’s unique traits and abilities (*63*). Early evidence indeed suggests that functional connectomes are associated with personality traits and even inter-personal closeness in social networks (*64, 65*).

Yet, the mechanistic implementation of these intriguing observations remains elusive. Inter-individual variability in the distribution of synaptic weights across the cerebrum, shaped through lifetime experiences according to Hebbian principles, may account — at least in part — for connectome fingerprinting (*63*). The heritability of the functional connectome has also been discussed, especially for fronto-parietal networks (i.e., dorsal and ventral attention network and the default mode network) (*66*–*68*). Heritability of brain spectral characteristics is also actively discussed (*69*–*71*). This emerging literature and the empirical evidence of brain fingerprinting certainly motivates more research on new, fascinating questions about the biological nature of the self.

### Sampling population diversity for personalized interventions

Robust individual signatures of brain activity may be transformative to neurophysiological phenotyping and population neuroscience. With the increasing availability of multi-omic data repositories, there is a research opportunity to span the diversity of statistical normative characteristics of brain fingerprints across the population in relation to behaviour, environmental and clinical variables (*1, 3, 27*). Our study highlights the utility of datasets of individuals who have been scanned on multiple occasions to capture and characterize interindividual variability as meaningful information. Ideally, large databanks of individual variants sampled across multiple dimensions of socio-economic, age, and geographic factors enable normative modeling approaches to establish the risk traits of developing syndromes of e.g., early cognitive decline, neurodegeneration or mental illness. Previous work has shown that mental disorders may affect the stability of individual fingerprints over time and therefore points at possible translational applications of the approach (*15, 72*). We may also foresee that changes over time or lack thereof of a person’s brain fingerprint may also constitute a new class of non-invasive markers of responses to neurological and other treatment of a variety of chronic, neurodegenerative or acute (e.g., stroke) conditions. Brain fingerprints derived from relatively short, task-free sessions may play a leading role to realize this vision in practice.

Brain fingerprinting may also contribute to future endeavours in establishing how oscillatory dynamics at rest support cognitive functions across the lifespan. MEG brain fingerprinting presents several potential advantages in terms of safety, shorter scan time, and immediate proximity of a care person during data collection, especially for special populations. The methodological approaches proposed herein can, in principle, transfer to EEG fingerprinting (*17*–*19*), which would be more readily available in clinics. Whether results would be as robust with EEG than with MEG remains to be demonstrated. Indeed, EEG source mapping is more prone to contamination from muscle artifacts and is more sensitive to approximations in the biophysical modeling of head tissues, which may compromise further fingerprinting capabilities (*27*).

In sum, our study extends the concept of neural or brain fingerprint to fast and large-scale resting-state electrophysiological dynamics, which encapsulate meaningful individual differences in both functional connectivity and neuroanatomical maps of power spectrum characteristics. We are hopeful that the present contribution paves the way to replication and extension using larger open datasets. Many fascinating outstanding questions remain about the biological nature of inter-individual variability expressed via neural oscillations and brain network dynamics, and more specifically how these differences associate with behavior and diseases natural history. The research ahead is for future population neuroscience studies.

## Material and Methods

### The Open MEG Archives (OMEGA)

We used data from the Open MEG Archives (OMEGA; 6) consisting of resting-state MEG recordings acquired using the same MEG system (275 channels whole-head CTF; Port Coquitlam, British Columbia, Canada). The sampling rate was 2400 Hz, with an antialiasing filter applied at 600 Hz cut-off, and built-in third-order spatial gradient noise cancellation (see 6 for details on data acquisition).

We analysed MEG resting-state data from 158 unrelated OMEG participants (77 Females, 31.9 ± 14.7 years old). Recordings were approximately 5-min long. Supplementary Table 1 provides details on scanning procedures and Supplementary Table 2 on demographics. A subset of these individuals (N=47) had recordings over multiple visits (different days) and were used in the *between-session* fingerprinting challenge. The OMEGA data management protocol was approved by the research ethics board of the Montreal Neurological Institute.

### MEG data preprocessing and feature extraction

MEG data were preprocessed using Brainstorm (73; version Oct-12-2018) (following good-practice guidelines (*74*). Unless specified, all steps below were performed using the Brainstorm toolkit, with default parameters. Line noise artifact (60 Hz) along with its 10 harmonics were removed using a notch filter bank. Slow-wave and DC-offset artifacts were removed using a high-pass FIR filter with a 0.3-Hz cut-off. We derived Signal-Space Projections (SSPs) to remove cardiac and ocular artifacts. We used electro-cardiogram and -oculogram recordings to define signal projectors around identified artifact occurrences. We also applied SSPs to attenuate low-frequency (1-7 Hz) and high-frequency noisy components (40-400Hz) due to saccades and muscle activity, respectively. Bandpass filtered duplicates of the cleaned data were produced for each frequency band of interest (delta: 1-4Hz, theta: 4-8Hz, alpha: 8-13Hz, beta: 13-30Hz, gamma: 30-50Hz, and high gamma: 50-150Hz). Distinct brain source models were then derived for all narrowband versions of the MEG sensor data.

Each individual T1-weighted MRI data was automatically segmented and labelled with Freesurfer (*75*). Coregistration with MEG sensor locations was derived using dozens of digitized head points collected at each MEG session. We produced MEG forward head models for each participant using the overlapping spheres approach, and cortical source models with LCMV beamforming, all using Brainstorm with default parameters (2016 version for source estimation processes). We performed data covariance regularization. To reduce the effect of variable source depth, the estimated source variance was normalized by the noise covariance matrix. Elementary MEG source orientations were constrained normal to the surface at 15,000 locations of the cortex. Noise statistics for source modeling were estimated from two-minute empty-room recordings collected as close as possible in time to each participant’s MEG session. Source timeseries were clustered into 68 cortical regions of interest (ROIs) defined from the Desikan-Killiany atlas (*37*) and dimension-reduced via the first principal component of all signals within each ROI. Connectome and spectral identification features were computed from ROI source timeseries. Individual functional connectomes were derived in all frequency bands from the amplitude envelope correlation (AEC) approach (*76*). ROI timeseries were Hilbert transformed and all possible pairs of resulting amplitude envelopes were used to derive the corresponding Pearson correlation coefficients, yielding a 68×68 symmetric connectome array. We used Welch’s method to derive power spectrum density (PSD) estimates for each ROI (*77*), using time windows of 2 seconds with 50% overlap sled over all ROI timeseries and averaged across all PSDs within each ROI. The resulting frequency range of PSDs was 0-150Hz, with a frequency resolution of 0.5 Hz.

### Code Availability

The connectome and spectral features were then exported to Python (3.7.6) for subsequent fingerprinting analyses. All codes for including preprocessing and data analysis can be found on the project’s GitHub (LINK).

### Data Availability

The power spectra and connectomes derived from the preprocessed OMEGA samples and used to identify individuals in the present study are available upon request from corresponding authors.

### Fingerprinting and self-identifiability

We used a fingerprinting approach directly adapted from fMRI connectome fingerprinting methods (*12, 14*), which relies on correlational scoring of individuals between datasets. A given *probe* participant is identified from a cohort by computing all Pearson correlation coefficients between the spectral or connectome features of said probe at one timepoint (e.g., *dataset 1*) and the entire cohort at a different timepoint (e.g., *dataset 2*). The entry presenting the highest correlation to the probe determined the probe’s estimated identity i.e., identified entry in the cohort. This approach is applied between all pairs of participants in the cohort, yielding an asymmetric correlation matrix spanning the cohort. We report scores of *identification accuracy as* the ratio between the number individuals correctly identified with the described procedure and the total number of individuals in the cohort. Identification accuracy scores are obtained from identification challenges from dataset 1 to dataset 2 and vice-versa, *within-* and *between-sessions*. Figure 1 details the definition of the dataset labels used, and Supplemental Material contains the results from across all combinations of datasets/sessions.

Amico and Goñi (2018) proposed an identifiability score to quantify, for a given participant, the reliability of its identification from others in the cohort. Here, we extend this notion with the introduction of a *self-identifiability* measure, **I**_**self**_. Let **A** be the correlation matrix spanning the cohort (square, asymmetric) between dataset 1 and dataset 2, and N be the number of participants to identify. We define **I**_**self**_ as the z-score of participant P_i_ ‘s correlation to themselves between dataset 1 and dataset 2, with respect to P_i_’s correlation to all other individuals in the cohort, noted: **I**_**self (i)**_ **= (**Corr_ii_ – μ_**ij**_) / σ_ij_, where Corr_ii_ is the P_i_’s correlation between dataset 1 and dataset 2, μ_ij_ is the mean correlation between participant P_**i**_ in dataset 1 and all other individuals in dataset 2 (i.e. the mean along the i^th^ row of matrix **A**), and σ_i_ is the empirical standard deviation of inter-individual features correlations. Thus, if a participant is easily identifiable, its self-identifiability increases; whereas small self-identifiability scores indicate a participant particularly difficult to identify from the rest of the cohort.

### Recording artifacts and self-identifiability

To investigate the effects of recording parameters and artifacts on fingerprinting, we related each individual’s self-identifiability to several possible confounds. The duration of each scan was compared to self-identifiability to verify that longer recordings available from a subset of individuals did not make them easier to identify. We also correlated the root mean square (RMS) of signals that measured ocular, cardiac, and head movement artifacts over the duration of the entire recording to participants’ self-identifiability score. For cardiac artifacts for instance, we derived the RMS of ECG recordings; for ocular artifacts we used the HEOG and VEOG electrode recordings; and for motion artifact we extracted the RMS of all three head coil signals that measured 3-D head movements. These derivations were conducted for both the connectome and spectral broadband *within-session* fingerprinting challenge.

### Fingerprinting across frequency bands

We replicated the above fingerprinting approach using data restricted to each frequency band of interest (delta 1-4Hz, theta 4-8Hz, alpha 8-13Hz, beta 13-30Hz, gamma 30-50Hz, and high gamma 50-150Hz). We report the identification accuracy obtained from each narrowband signal in both the spectral and connectome fingerprinting challenges in Figure 2 and Figure 3, for the *within-* and *between-session* fingerprinting challenges respectively. We also performed fingerprinting tests based on sensor data only. We used the same connectome and spectral approaches as the MEG source maps, considering the time series of each of the 275 MEG channels instead of the 68 ROI time series derived from the brain map parcels. We report the identification performances from both the sensor and source analyses in Figure 3 and in Supplemental Material.

### Between-session and shortened fingerprinting challenges

We verified the robustness of MEG fingerprinting with respect to 1) the ability to identify participants over time and 2) from reduced data durations. We subdivided participants into three additional challenges: the *within-session—shortened, between-session*, and *between-session—shortened* challenge. First, we used the participant data described in the *within-session* analysis and extracted connectome and spectral fingerprinting features over three 30-second non-overlapping time segments. This duration was based on the length of the shortest recording in the data sample (Figure 1aii). We applied the same fingerprinting procedure as described in **Fingerprinting and self-identifiability** across all possible combinations of the three 30-second datasets. Second, we assessed the stability of the fingerprinting outcomes using a subset of participants with consecutive MEG sessions separated by several days (N=47; separated on average by 201.7 days, see Supplemental Materials for details). Again, we applied the same fingerprinting procedure as described in **Fingerprinting and self-identifiability** for this *between-session* challenge. Lastly, we applied the same shortened analysis—described above—to the subset of individuals with multiple scans (i.e., the between-sessions data). We report all possible combinations of datasets (i.e., three 30s segments from day 1 and three 30s segments from day 2; see Figure 1a for example) in Figure 3.

### Empty-room fingerprinting

We tested whether environment and instrument noise daily conditions would bias individual identification using empty-room recordings collected from each MEG session. The empty-room data was processed identically to the participants data, using the same individual imaging kernels, and were used to identify participants. We ran all possible combinations of empty-room vs. participants datasets (e.g., empty-room 1 vs. participant dataset 1, empty-room 2 vs. participant dataset 1, etc.) and computed the sample mean of the identification accuracies across all dataset combinations. The identification accuracies obtained represent estimates of baseline reference performances that can be compared to each form of fingerprinting based on actual participant data (i.e., connectome or spectral, broadband or band-specific; see Figure 2 and Supplemental Material). In a similar fashion, we also used sensor-level empty-room recordings of each participant for fingerprinting—attempting to identify individuals’ recordings from their empty-room features. The results of this analysis are reported in the Supplemental Material.

### Most characteristic features for fingerprinting

We quantified the contribution of each feature (i.e., edges in the connectivity matrix or a frequency band in an anatomical parcel) towards identifying individuals using Intraclass Correlations (ICC). ICC is commonly used to measure the agreement between two observers (e.g., ratings vs. scores). The stronger the agreement, the higher the ICC (*12, 38*). ICC derives a random effects model whereby each item is rated by different raters from a pool of potential raters. We selected this measure to capture the inter-rater reliability of each participant as their own rater to identify which edges (e.g., connections in FC) are the most consistent (i.e., which features of a participant P_i_ in dataset 1 are most like dataset 2). Here, the higher the ICC, the more consistent a given feature was within individuals. Additionally, we computed two other measures of edgewise contribution proposed by Finn and colleagues (*14*): *group consistency* and *differential power* (Supplemental Material). We applied all measures (i.e., ICC, group consistency, and differential power) in the context of the broadband *within-session* fingerprinting challenge. The source maps shown in Figure 4, Figure 5 and Supplemental material were generated using R (V 3.6.3; *74*) with the *ggseg* package (*79*).

### Partial Least-Squares: MEG features of participant demographics

We conducted a Partial Least-Squares (PLS) analysis with the Rotman-Baycrest PLS toolbox (*80*). PLS is a multivariate statistical method that relates two matrices of variables (e.g., neural activity and participant demographics) by estimating a weighted linear combination of variables from both data matrices to maximize their covariance. The associated weights can be interpreted neural patterns (e.g., functional connections) and their associated demographic profiles. PLS used singular value decompositions of the z-scored neural activity-demographics covariance matrix. This decomposition yielded orthogonal latent variables (LV) associated to a pattern of neural activity (i.e., functional connectivity or spectral power) and demographics. To assess the significance of these multivariate patterns, we computed permutation tests (10,000 permutations). Each permutation shuffled the order of the observations (i.e., the rows) of the demographic data matrix before running PLS on the resulting surrogate data under the null hypothesis that there was no relationship between the demographic and neural data. A *p*-value for the LVs was computed as the proportion of times the permuted singular values exceeded that of the original data. We explored the first significant LV from the broadband connectome and spectral fingerprinting features. We also assessed the contribution of each variable in the demographics and neural activity matrices by bootstrapping observations with replacement (10,000 bootstraps). We computed 95-% confidence intervals for the demographic weights and bootstrap ratios for the neural weights. The bootstrap ratio was computed as the ratio between each variable’s weight and the bootstrap-estimated standard error.

## Author Contribution

All authors conceptualized the study, J.D.S.C. and H.D.O. preformed the analyses, S.B. and B.M. provided guidance with data interpretation, J.D.S.C. wrote the first draft of the manuscript, all authors contributed to the writing and editing of the manuscript.

## Supplemental material

### MEG fingerprinting is robust against sample demographics

The OMEGA data repository contains 158 participants, with a subset (N=47) scanned at multiple occasions several days apart. OMEGA consists essentially of data from healthy controls with a 18-73-year age span (SD=14.7 years; Supplemental Table 1).

One potential confound that could have inflated our ability to fingerprint individuals is the heterogeneity introduced by both healthy and clinical populations in the OMEGA cohort. To address this concern, we ran a secondary analysis where we performed the fingerprinting procedures described in the manuscript with only healthy controls (N=130). The results, reported in Supplemental Table 2, demonstrated that fingerprint performances were not biased by the patients/controls heterogeneity of the OMEGA sample. We observed a decrease of less than 1% in performance relative to fingerprinting from the entire cohort. Further, there was no clear relationship between self-identifiability and demographics (Figure S1)., using connectome (age: r= 0.08, p = 0.2; gender: t= -0.27, p = 0.7; handedness: t= -0.51, p = 0.6; clinical status: t= -0.87, p = 0.3; two-tailed) and spectral fingerprinting (age: r= 0.10, p = 0.1; gender: t= 0.62, p = 0.5; handedness: t= 0.13, p = 0.8; clinical status: t= 0.84, p = 0.3; two-tailed).

**Figure S1:**
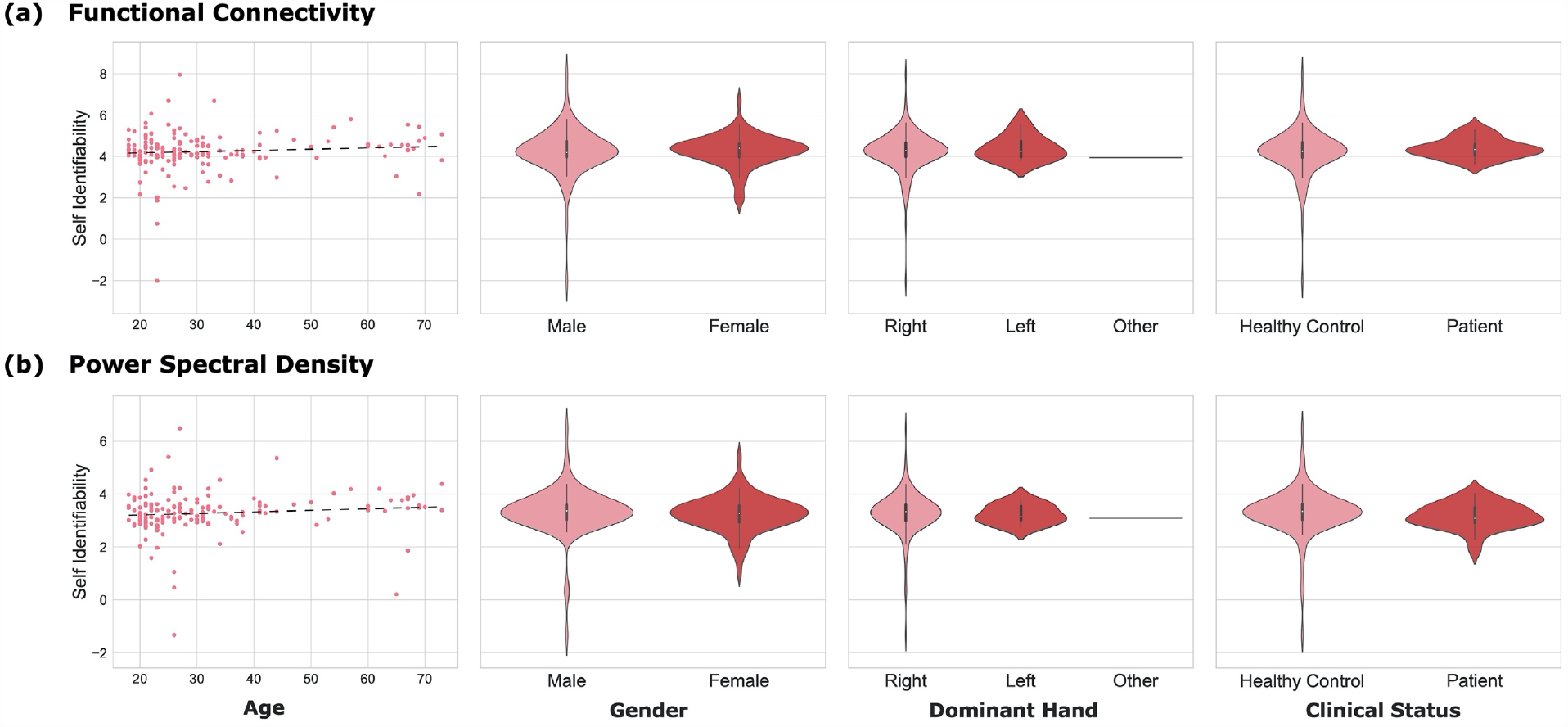
Self identifiability is not associated with demographics. The plots depict demographic variables and corresponding self-identifiability scores across both **(a)** connectome and **(b)** spectral broadband *within-session* fingerprinting. Demographic variables included age, biological sex, dominant hand, and healthy vs. patient categories. There was no clear relationship between demographics and self-identifiability — i.e., differences in demographics did not drive self-identifiability.

Acquisition parameters did not affect both fingerprinting performances (Figure S2). Participants with longer recordings (i.e., more data) were not more identifiable (connectome: r= -0.02, p = 0.7; spectral: r= 0.02, p = 0.8). This observation is consistent with the *within- & between-session shortened* fingerprinting results, which demonstrate individuals were identifiable from shorter 30-second recordings (see below).

Taken together, these supplemental results demonstrate that MEG fingerprinting is robust against data artifacts, heterogeneous sample demographics and acquisition parameters.

**Supplemental table 1:** OMEGA participant demographics. Demographic variables summarized for both subsets of the OMEGA data repository.

|  | Within-session data | Between-session data |
| --- | --- | --- |
| Age | 31.9 ± 14.7 | 26.7 ± 11.6 |
| Gender | 77 Females | 24 Females |
| Dominant Hand | 147 Right, 8 Left, 1 Other | 44 Right, 3 Left |
| Clinical Status | 130 Healthy Controls<br>22 ADHD<br>6 Chronic Pain | 25 Healthy Controls<br>22 ADHD |

**Figure S2:**
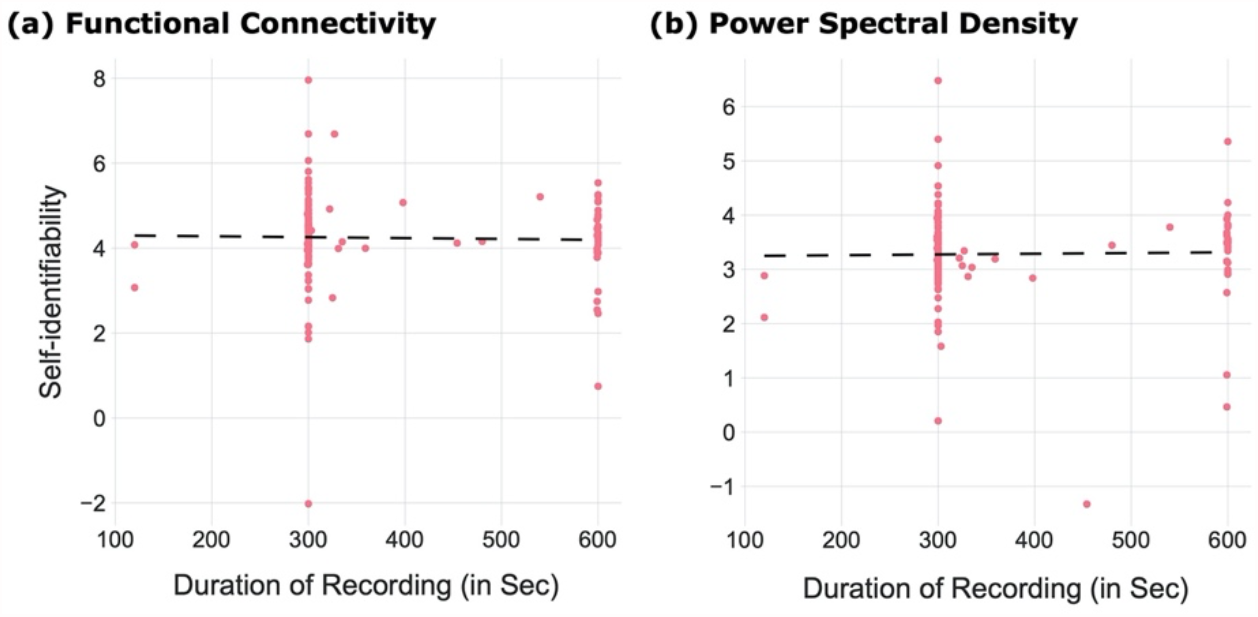
Recording duration did not affect self-identifiability. Scatter plots of self-identifiability vs. duration of data collections, for the broadband *within-session* challenge. There was no clear relationship between self-identifiability and the duration of the MEG recordings across participants.

**Supplemental table 2.** Fingerprinting performances of healthy controls. Identification performances of connectome and spectral broadband *within-session* fingerprinting obtained from for the entire repository (healthy controls and patients), and from healthy participants only. Each column reports fingerprinting performances from Dataset 1 to Dataset 2 and vice-versa (see Figure 1 for details). Overall, identification accuracy decreased slightly by ∼0.9% when comprising healthy participants only. Consistent with our findings reported in Figure S2, clinical status did not play a major role in the identification of individuals.

|  | All Participants |  | Only Healthy Controls |  |
| --- | --- | --- | --- | --- |
|  | Dataset 1 to Dataset 2 | Dataset 2 to Dataset 1 | Dataset 1 to Dataset 2 | Dataset 2 to Dataset 1 |
| Connectome | 94.9% | 94.3% | 93.8% | 93.0% |
| Spectral | 96.2% | 96.2% | 95.3% | 95.3% |

**Figure S3:**
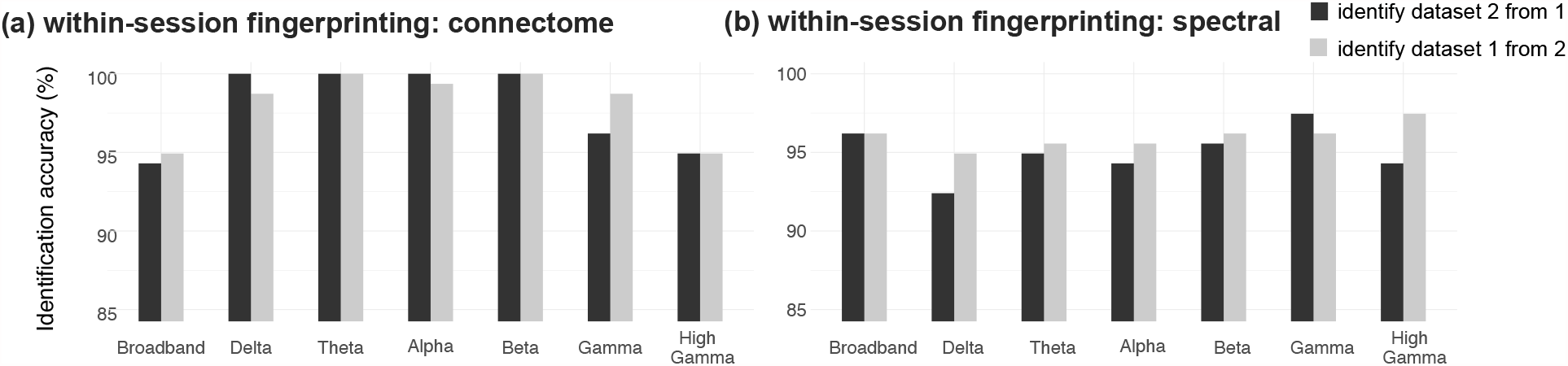
Identification accuracy from within-session datasets. Results from MEG *within-session* fingerprinting. Identification accuracy for **(a)** connectome and **(b)** spectral fingerprinting (broadband and narrowband data). The accuracy scores are reported for identification from dataset 1 to dataset 2 and vice-versa, as explained in Methods.

**Figure S4:**
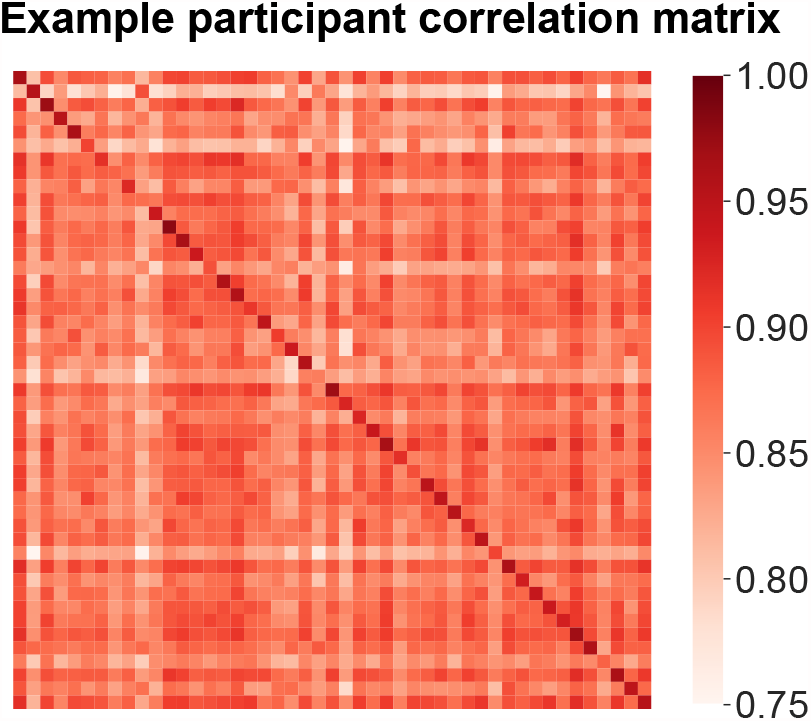
Example participant correlation matrix for fingerprinting. Exemplar participant correlation matrix derived from *between-session* data used for fingerprinting. The study-identity of participants was determined by the highest correlation statistics taken across rows (e.g., to identify dataset-2 from dataset-1) or columns (to identify dataset-1 from dataset-2).

### Data reduction from principal component analysis does not improve MEG fingerprinting substantially

Amico and Goñi (*1*) previously reported improvements to participant differentiation when using data reduction techniques prior to identification, using e.g., principal component analysis (PCA). We reproduced their approach, using PCA to reduce the dimensionality of the connectome and spectral feature spaces prior to fingerprinting. Our results provided little support to PCA reconstruction improving identification accuracy, as shown Figure S5 and in Supplemental Table 3. PCA increased self-identifiability by less than 1.5%. Data reduction had limited beneficial impact possibly because of high fingerprinting performances at baseline (without data reduction). We also emphasize that we conducted MEG source time series extraction via a PCA of all local time series within each parcel. It is therefore likely that this dimension reduction procedure contributed to improve signal-to-noise ratio and limited the impact of subsequent PCA of features.

**Supplemental Table 3:** Limited contribution of data reduction from principal component analysis to MEG fingerprinting. Performances in identification accuracy for connectome and spectral broadband *within-session* fingerprinting, for both original and PCA-reconstructed data (*1*). PCA data reduction improved connectome fingerprinting performances only slightly (about 2%). It had virtually no effect on spectral fingerprinting performances.

|  | Original<br>(un-reconstructed) |  | PCA<br>Reconstructed |  |
| --- | --- | --- | --- | --- |
|  | Dataset 1 to<br>Dataset 2 | Dataset 2 to<br>Dataset 1 | Dataset 1 to<br>Dataset 2 | Dataset 2 to<br>Dataset 1 |
| Connectome | 94.9% | 94.3% | 96.2% | 96.2% |
| Spectral | 96.2% | 96.2% | 96.2% | 96.2% |

**Figure S5:**
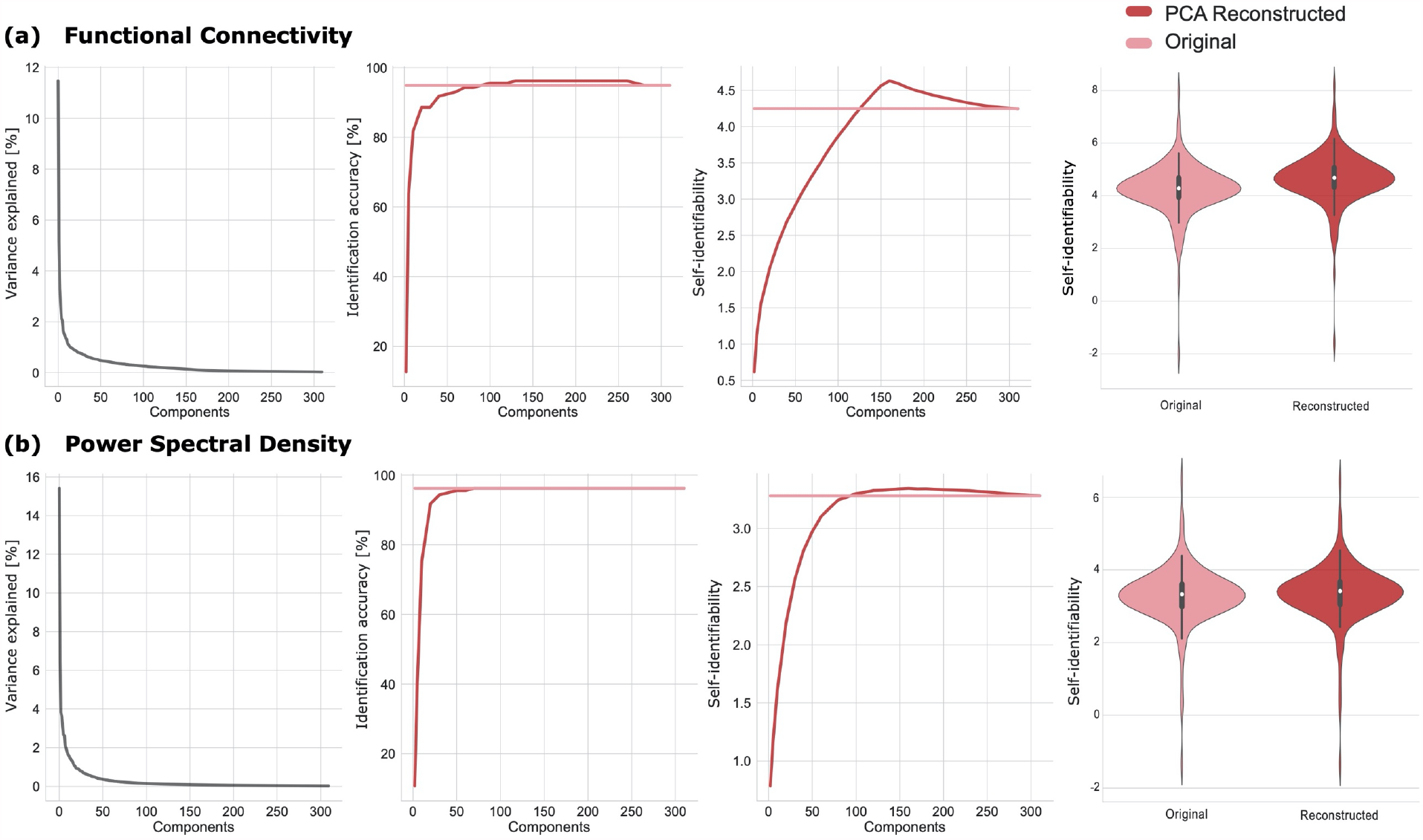
Limited benefit of PCA reconstruction to identification accuracy. PCA reconstruction as proposed by Amico and Goñi (2018) had limited effect on **(a)** connectome and **(b)** spectral *within-session* fingerprinting. The original results (Figure 2) are plotted against PCA-reconstructed results. From left to right, plots show *i)* PCA components plotted vs. their respective fractions of signal variance explained, *ii)* identification accuracy across PCA components, *iii)* average self-identifiability across PCA components, and *iv)* violin plots of self-identifiability before and after PCA reconstruction. Overall, PCA reconstruction did not substantially improve identification accuracy.

### Fingerprinting with 30-second data segments

We challenged MEG fingerprinting using short 30-second data segments (i.e., shortened *within-session* fingerprinting). We epoched participants’ MEG recordings into three datasets of 30 second, where the first dataset was the first 30 seconds of the recording after having removed the initial five seconds, the second dataset was the 30 seconds immediately following the first dataset, and the last dataset was the last 30-second segment of the recording after having removed the last ten seconds (Figure 1). Cropping the initial and last few seconds from recordings excluded edge filtering and other session artifacts. The lengths of the short datasets and epochs were determined from the participant with the shortest available recording. This procedure yielded three data segments for fingerprinting purposes via 6 possible dataset pairs (i.e., dataset 1 and 2; dataset 2 and 3; and dataset 1 and 3 and vice-versa). Results for all possible combinations of datasets are reported in Figure S6.

Connectome fingerprinting successfully identified individuals across all possible combinations of datasets (Figure S6). Identification from recordings collected closer in time (e.g., dataset-1 and dataset-2) outperformed identification from datasets collected further apart in time (e.g., between dataset-1 and dataset-3). Overall, spectral fingerprinting yielded lower identification accuracy than connectome fingerprinting, in particular from datasets further apart in time.

In a similar fashion, we challenged MEG fingerprinting using short 30-second data segments from different sessions (i.e., *between-session* fingerprinting). This yielded 6 epochs of data for fingerprinting (i.e., three from both the first and second recording, see Figure 1a). Identification results averaged across all possible data pairs are reported Figure 3c. Connectome fingerprinting performances were greater than those from spectral fingerprinting. Identification from slower frequency data components performed worse in comparison to higher bands – see main article body for discussion.

**Figure S6:**
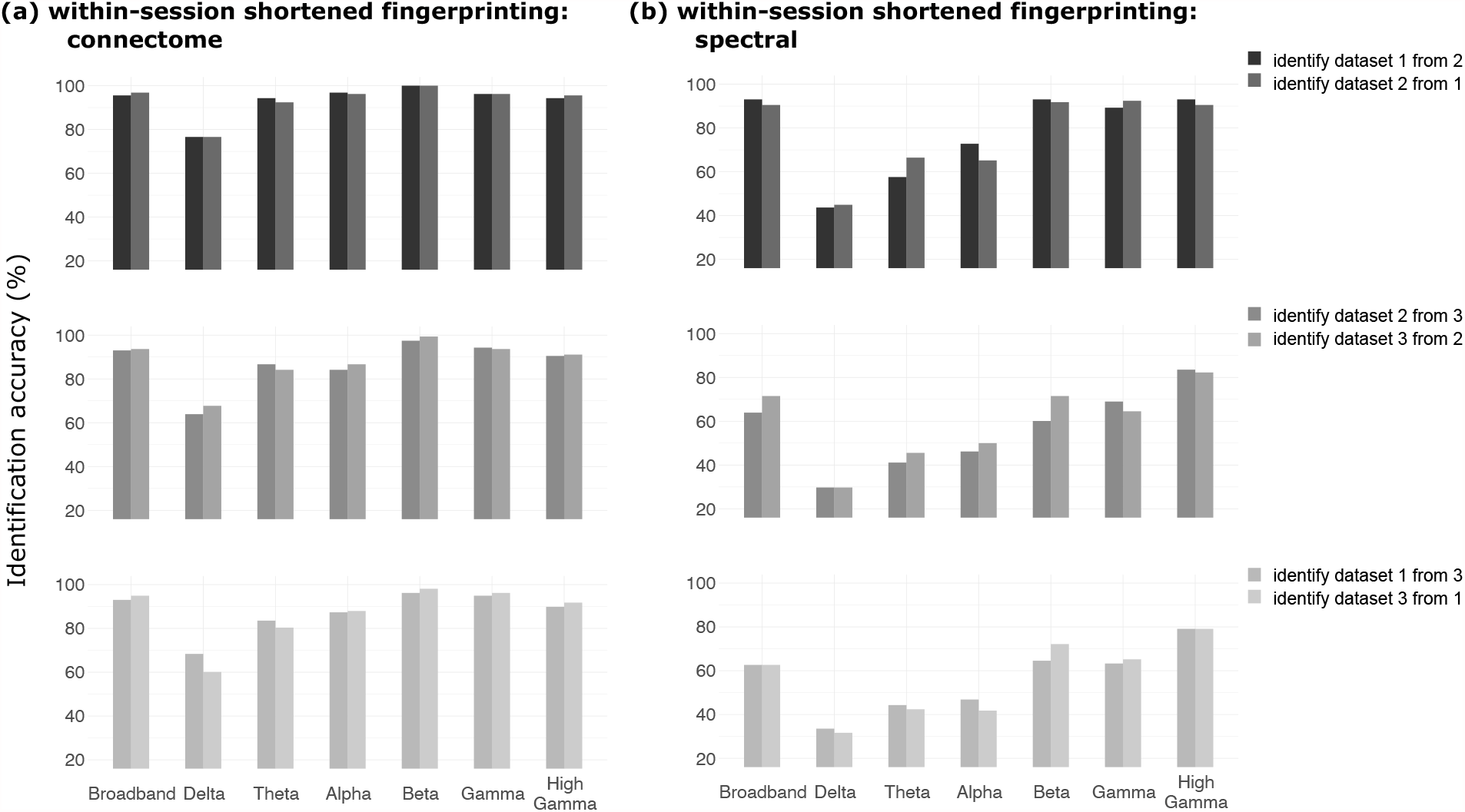
Identification accuracy from shortened within-session datasets. Identification results from shortened *within-session* datasets (30 seconds) for **(a)** connectome and **(b)** spectral broadband and narrowband fingerprinting. The accuracy scored are reported for identification from all possible combinations of datasets, (i.e., dataset 1 to predict dataset 2, dataset 3 to predict dataset 2, etc.; see Methods for details). Identification accuracy increased as datasets were proximal in time (i.e., fingerprinting accuracy for dataset 1 to dataset 2 was greater than for dataset 1 to dataset 3).

### Fingerprinting across recording sessions

We also report fingerprinting accuracy performances from all possible pairs of datasets for the *between-session* fingerprinting challenge in Figure S7. Overall, spectral fingerprinting outperformed connectome fingerprinting, as discussed in the main text.

**Figure S7:**
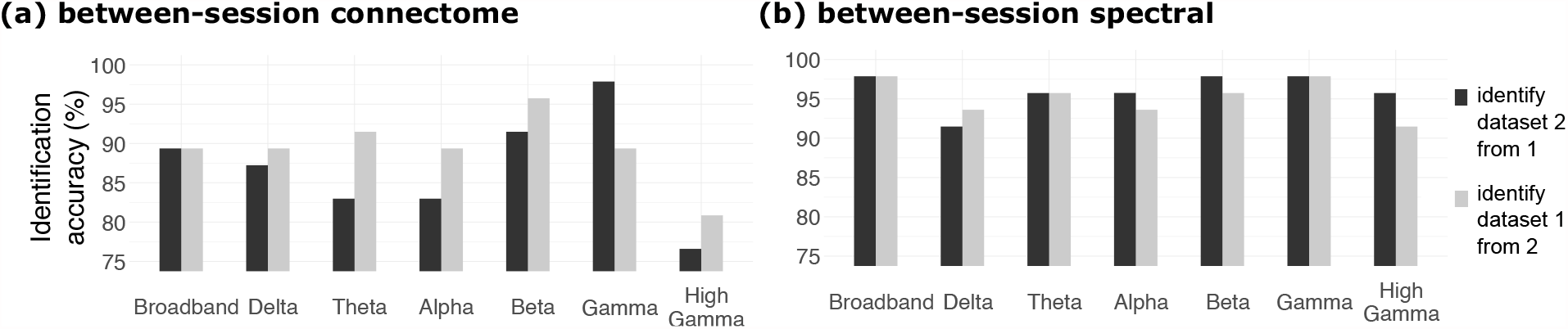
Between-session identification accuracy. Results from MEG *between-session* fingerprinting. Identification accuracy for both **(a) c**onnectome and **(b)** spectral broadband and narrowband fingerprinting. The accuracy scores are reported for identification from dataset 1 to dataset 2 and vice-versa (see Methods).

### Individuals cannot be identified from their respective imaging kernels

We verified that the within-session identification of individuals was not possible from empty-room data (i.e., with no participant under the MEG sensor array) processed through their respective imaging kernel of beamformer weights. Indeed, these latter are defined from individual anatomy and head position under the MEG sensor array, which may have been sufficient information to drive identification. We therefore ran the same fingerprinting pipeline on each session’s empty-room data transformed through the corresponding individual’s beamformer imaging kernel, which was identical for each of the within-session data segments used. Note that for the between-session challenges, the imaging kernels were adjusted to the respective individual head positions measured during each session. These analyses demonstrated that the imaging kernel information did not contribute substantially to MEG fingerprinting (overall performance was below 20% on average, See Figure 2).

We also ran the MEG fingerprinting pipeline directly from the sensor data of the empty-room recordings, without transformation through individual imaging kernels, to assess the floor level of identification performances from non-brain data only. The data confirmed substantially lower levels of identification (<5% accuracy on average; see Figure S8).

**Figure S8:**
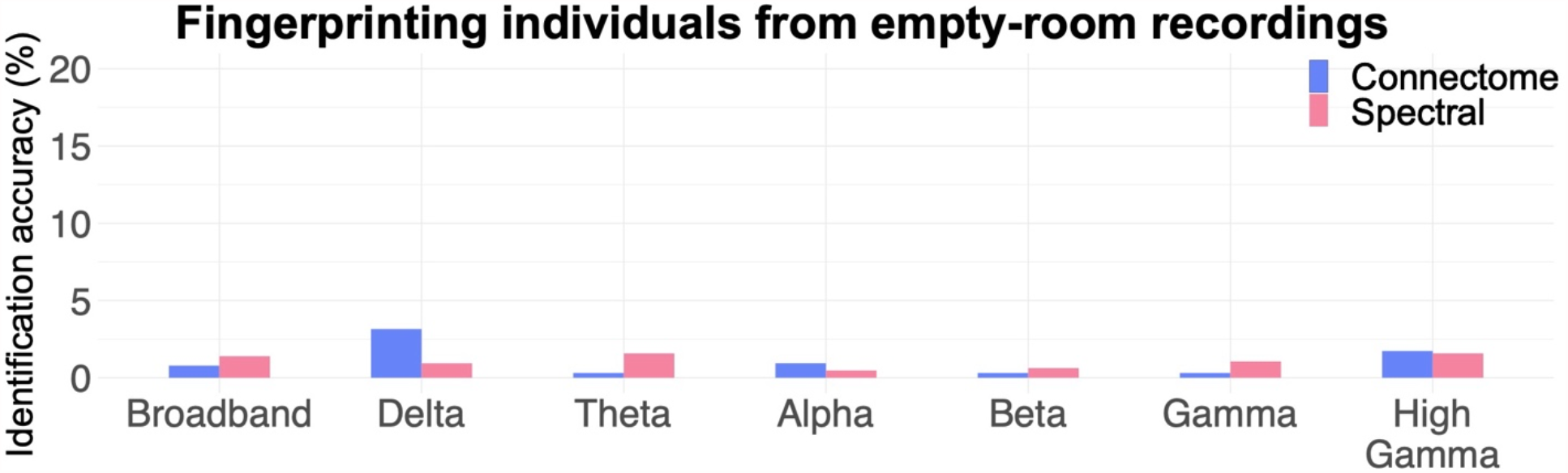
Verification of failed fingerprinting from non-brain data (empty-room recordings) Results for the empty-room sensor fingerprinting challenge. As expected, identification accuracies of connectome and spectral broadband and narrowband fingerprinting were substantially lower than from actual MEG data with individuals present.

### Fingerprinting from scalp data only

We also performed MEG fingerprinting from individual sensor data, with no MEG source reconstruction to assess the added value of source modeling. We replicated the above MEG fingerprinting pipelines from the *within*-, *within-shortened*, and *between-session* analyses. Identification performances were less than with source modeling, especially from signal components in higher frequency bands and for the *shortened* challenges (see Figure S9, S10, & S11). Yet for other signal components and longer durations, individuals remain identifiable from sensor-level data collected between sessions (>60% accuracy from broadband data), albeit with lower accuracy than when using MEG source transformations, which explicitly account for different head positions between sessions.

Taken together with the empty-room fingerprinting tests above, these results provide evidence that brain signals, not environmental conditions, were crucial for individual identification.

**Figure S9:**
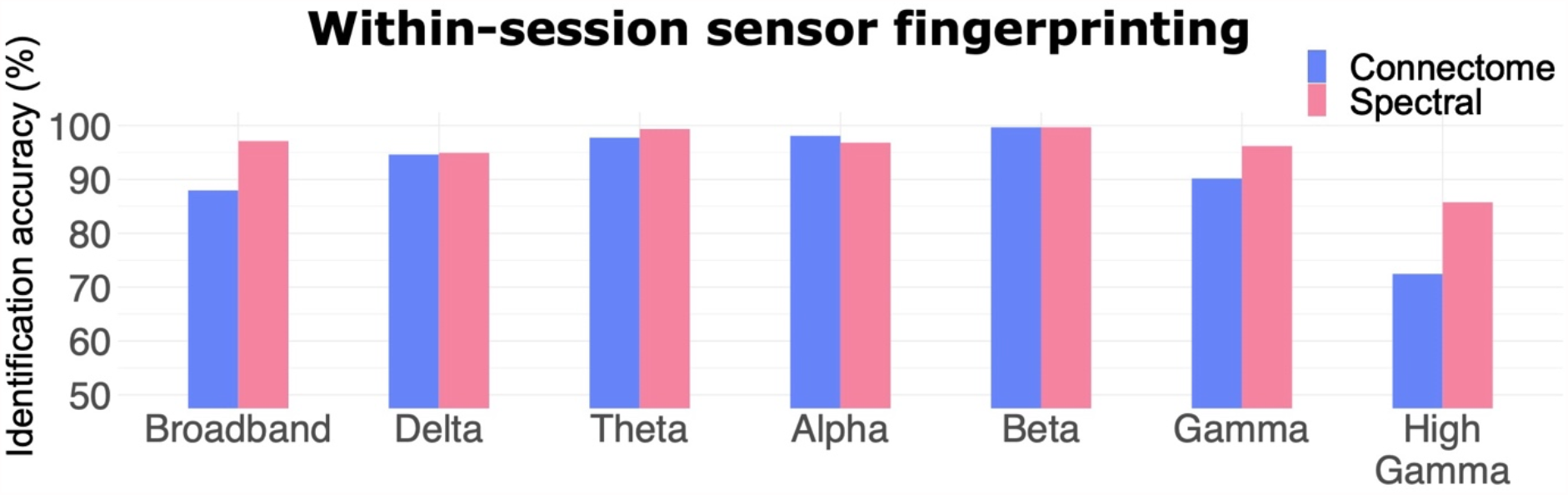
Within-session identification from MEG sensor data (no source modeling) Results from MEG sensor data in the *within-session* fingerprinting challenge. The identification accuracy statistics are shown for both connectome and spectral broadband and narrowband fingerprinting. The average accuracy scores are reported across identifications from dataset-1 to dataset-2 and vice-versa (see Methods).

**Figure S10.**
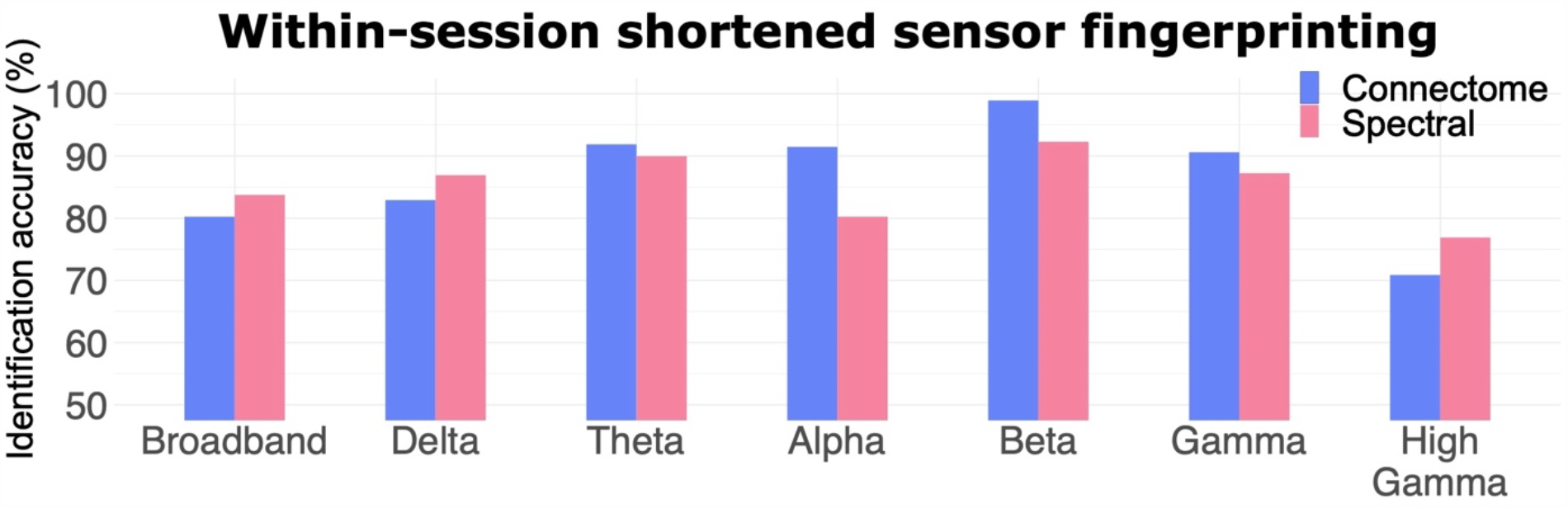
Within-session identification from shortened (30-s) MEG sensor data (no source modeling) Results from MEG sensor data in the *within-session shortened* fingerprinting challenge. The identification accuracy statistics are shown for both connectome and spectral broadband and narrowband fingerprinting. The average accuracy scores are reported across identifications from all possible pairs of datasets (see Methods).

**Figure S11:**
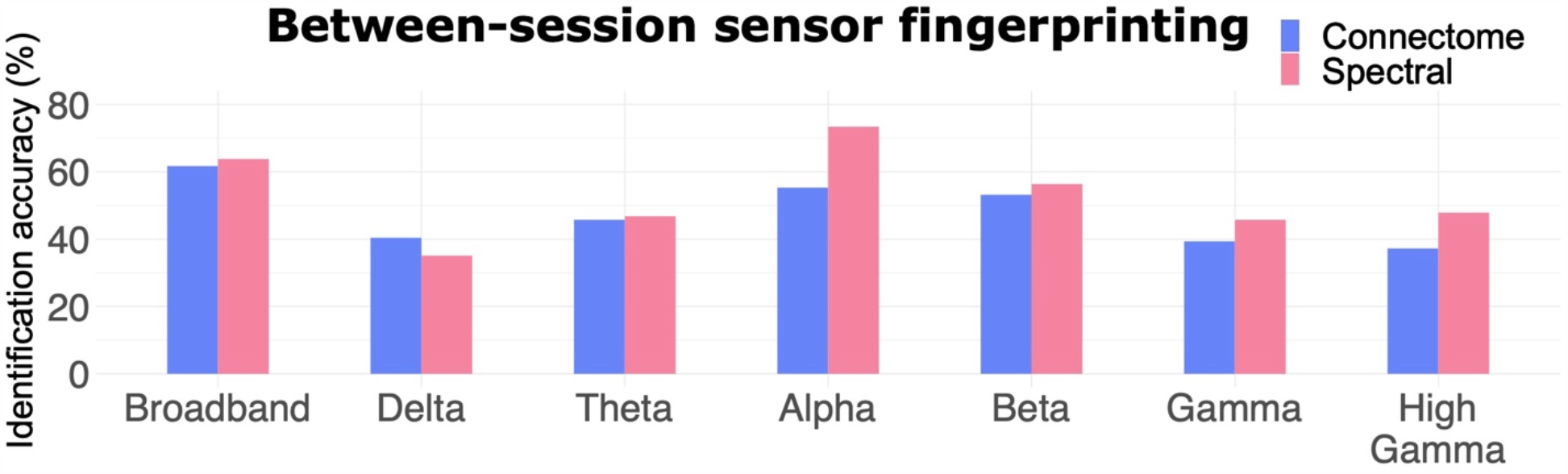
Between-session identification from MEG sensor data (no source modeling) Results from MEG sensor data in the *between-session* fingerprinting challenge. The identification accuracy statistics are shown for both connectome and spectral broadband and narrowband fingerprinting. The average accuracy scores are reported across identifications from dataset-1 to dataset-2 and vice-versa (see Methods).

### Salient neurophysiological features for fingerprinting

We reported in the main manuscript intraclass correlations (ICC) to determine which features contributed to individual identification the most. We also performed two additional analyses, deriving *group consistency* and *differential power*. These two metrics were proposed by Finn and colleagues (*2*) to identify the features which were the most consistent across their cohort, vs. The features which were the most consistent within individuals but different across participant, respectively (*2*). Differential power measures the empirical probability that a given feature is more likely to have a higher edgewise product vector across individuals than within the same individual. Taking the sum of the natural log of this probability across subjects yields differential power (*2*). The higher the differential power, the better a feature discriminates between individuals. Results for differential power are plotted in Figures S7 and S9. We found that the most discriminant connectome features were the visual and limbic networks across frequency bands, while the most discriminant spectral features remained along midline structures for fast oscillatory signal components. Overall, these results confirmed the ICC analysis results, with the addition of the contributions of spectral power in the beta and gamma band along the supplementary motor, motor, and somatosensory cortices.

**Figure S12:**
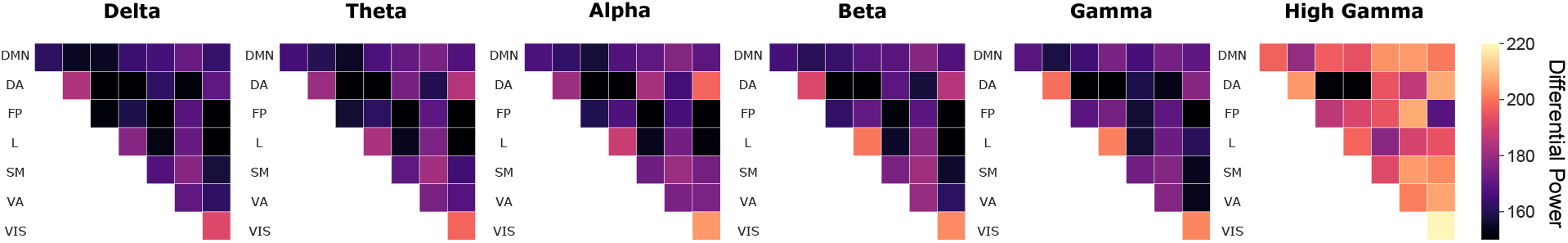
Differential power connectome fingerprinting. Differential Power (DP) analysis for broadband connectome fingerprinting of the *within-session* dataset (see Figure 1). Mean DP plotted within frequency bands and per resting-state network as defined by (*3*): Default Mode Network (DMN), Dorsal Attention (DA), Frontal-Parietal (FP), Limbic (L), Somato-Motor (SM), Ventral Attention (VA), and Visual (VIS). The higher the DP, the more the corresponding functional connection was essential for fingerprinting. The outstanding connections determined by DP for fingerprinting were the Visual network across all frequency bands, and the Limbic network in the beta and gamma bands.

Group consistency reflects edges that are consistent across individuals. Group consistency was computed from the mean edgewise product vector across all subjects (*2*). Large values of group consistency highlight features that are consistent both within participants and across the cohort. Our analyses are shown Figures S8 and S10. The resulting most consistent connectome features remained along the diagonal of the FC matrix (i.e., connections within the same networks) specifically in the Dorsal Attention and Fronto-Parietal networks. The most consistent features for spectral fingerprinting were in the lower frequency bands, specifically in the lateral frontal cortices. This outcome was consistent with our ICC results (see Manuscript).

**Figure S13:**
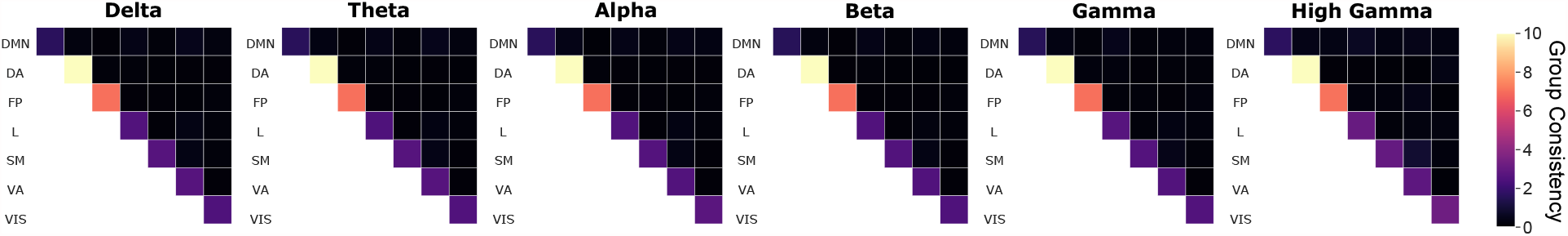
Group consistency connectome fingerprinting. Group Consistency (GC) analysis for broadband connectome fingerprinting of the *within-session* dataset (see Figure 1). Mean GC plotted within frequency bands according to the labels from (*3*): Default Mode Network (DMN), Dorsal Attention (DA), Frontal-Parietal (FP), Limbic (L), Somato-Motor (SM), Ventral Attention (VA), and Visual (VIS). The higher the GC, the more consistent was a functional connection within an individual and across the cohort. The most consistent connections were those along the diagonal, specifically for the Dorsal Attention and Frontal-Parietal networks across all frequency bands.

**Figure S14:**
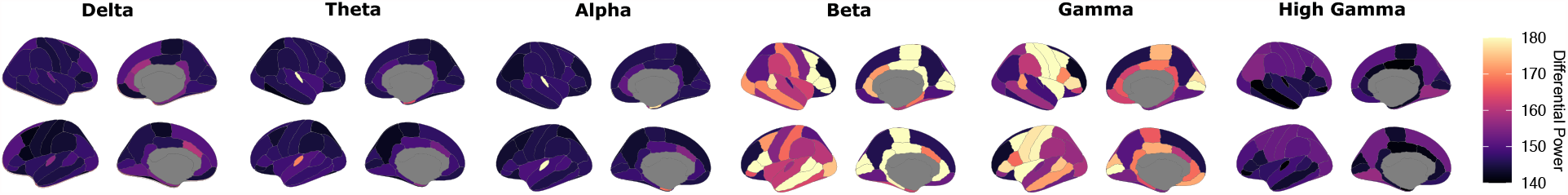
Differential power spectral fingerprinting. Differential Power (DP) analysis for broadband spectral fingerprinting of the *within-session* dataset (see Figure 1). Mean DP plotted within frequency bands according to the Desikan-Killiany atlas (*4*). The higher the DP, the more a given frequency band and ROI distinguished between individuals. The most characteristic regions and frequencies were medial structures for the beta band, and temporal and central regions for gamma band signals.

**Figure S15:**
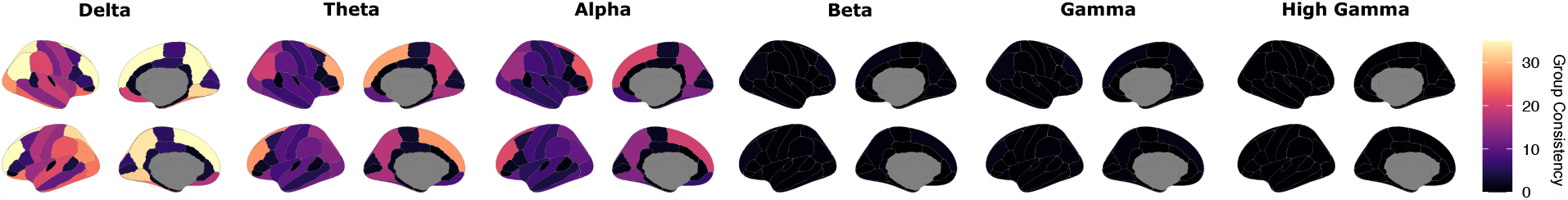
Group consistency spectral fingerprinting. Group Consistency (GC) analyses for broadband spectral fingerprinting of the *within recording session* dataset (see Figure 1). Mean GC plotted within frequency bands according to the Desikan-Killiany atlas (*4*). The higher the GC, the more a given frequency band and ROI remained consistent within individuals and across the cohort. The most stable frequencies were the lower bands (delta and theta) and the most consistent regions across individuals were lateral frontal areas.

### Partial Least Squares (PLS) analysis

We tested whether differences in resting-state neurophysiological signals related to meaningful demographic features using an exploratory Partial Least Squares (PLS) analysis. PLS is a multivariate statistical method that relates two data matrices based on latent variables (LV) that explain the highest covariance between the two datasets. Here, our two datasets consist of a demographic matrix (i.e., age, gender, handedness, and clinical status) and a neurophysiological data matrix (i.e., spectral power or functional connectome). Latent variables (which explain the most covariance between both matrices), and their corresponding variance explained are plotted in Figure S16. Significance of each latent variable was assessed via permutation tests. Permuting the rows of the data allowed us to compute an associate p-value for each latent variable (see Manuscript). We chose to explore the first significant latent variable which explained the most variance for each neurophysiological signal feature (i.e., the first component for connectomes and spectral data). The resulting weights associated to the latent neural and demographic components are depicted Figure 5 along with their bootstrapped ratios. These results corroborate how neurophysiological signals at rest, in addition to identifying individuals, carry meaningful information about participant demographics.

**Figure S16:**
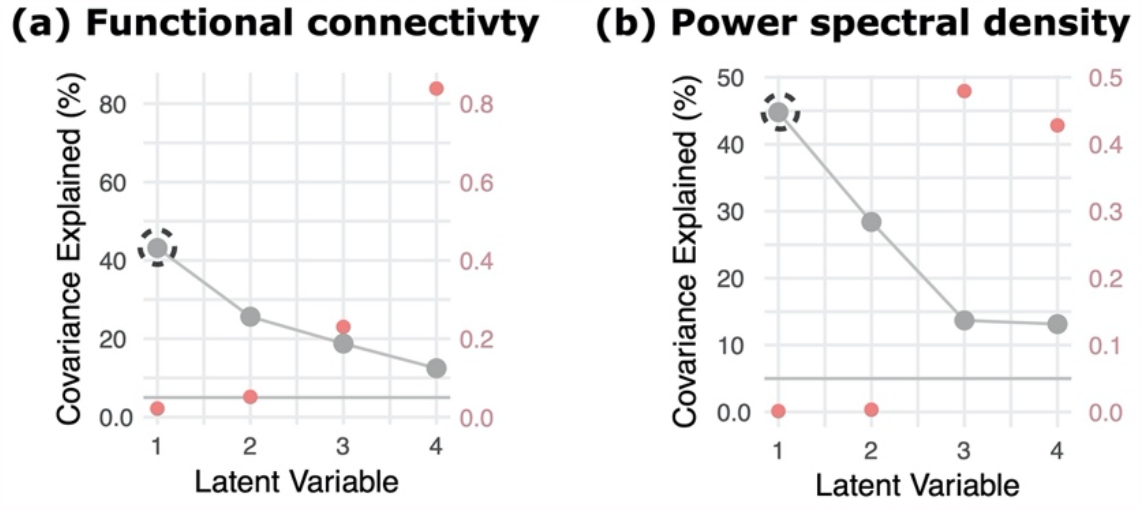
PLS latent variables. Results for the PLS analysis conducted for both **(a)** connectome and **(b)** spectral fingerprinting features. Each plot depicts the latent components obtained for each of the PLS analyses, their corresponding variance explained, and permuted *p*-value (right axis). One significant latent variable explained 43.1% of the variance for connectome fingerprinting and two latent variables explained 44.7% and 28.3% of the variance for spectral fingerprinting, respectively. We explored in the main Manuscript only the first significant component for each method (i.e., the circled component).

## References

1. J. Dubois, R. Adolphs, Building a Science of Individual Differences from fMRI. Trends Cogn. Sci. 20, 425–443 (2016).

2. M. B. Miller, J. D. Van Horn, Individual variability in brain activations associated with episodic retrieval: a role for large-scale databases. Int. J. Psychophysiol. Off. J. Int. Organ. Psychophysiol. 63, 205–213 (2007).

3. J. D. Van Horn, S. T. Grafton, M. B. Miller, Individual Variability in Brain Activity: A Nuisance or an Opportunity? Brain Imaging Behav. 2, 327 (2008).

4. T. Yarkoni, in APA handbook of personality and social psychology, Volume 4: Personality processes and individual differences., M. Mikulincer, P. R. Shaver, M. L. Cooper, R. J. Larsen, Eds. (American Psychological Association, Washington, 2015; http://content.apa.org/books/14343-002), xpp. 61–83.

5. D. S. Marcus, J. Harwell, T. Olsen, M. Hodge, M. F. Glasser, F. Prior, M. Jenkinson, T. Laumann, S. W. Curtiss, D. C. Van Essen, Informatics and data mining tools and strategies for the human connectome project. Front. Neuroinformatics. 5, 4 (2011).

6. G. Niso, C. Rogers, J. T. Moreau, L.-Y. Chen, C. Madjar, S. Das, E. Bock, F. Tadel, A. C. Evans, P. Jolicoeur, S. Baillet, OMEGA: The Open MEG Archive. NeuroImage. 124, 1182–1187 (2016).

7. R. A. Poldrack, K. J. Gorgolewski, Making big data open: data sharing in neuroimaging. Nat. Neurosci. 17, 1510–1517 (2014).

8. R. B. Mars, R. E. Passingham, S. Jbabdi, Connectivity Fingerprints: From Areal Descriptions to Abstract Spaces. Trends Cogn. Sci. 22, 1026–1037 (2018).

9. B. Mišić, O. Sporns, From regions to connections and networks: new bridges between brain and behavior. Curr. Opin. Neurobiol. 40, 1–7 (2016).

10. S. A. Valizadeh, F. Liem, S. Mérillat, J. Hänggi, L. Jäncke, Identification of individual subjects on the basis of their brain anatomical features. Sci. Rep. 8, 5611 (2018).

11. C. Wachinger, P. Golland, W. Kremen, B. Fischl, M. Reuter, BrainPrint: A discriminative characterization of brain morphology. NeuroImage. 109, 232–248 (2015).

12. E. Amico, J. Goñi, The quest for identifiability in human functional connectomes. Sci. Rep. 8, 8254 (2018).

13. S. Bari, E. Amico, N. Vike, T. M. Talavage, J. Goñi, Uncovering multi-site identifiability based on resting-state functional connectomes. NeuroImage. 202, 115967 (2019).

14. E. S. Finn, X. Shen, D. Scheinost, M. D. Rosenberg, J. Huang, M. M. Chun, X. Papademetris, R. T. Constable, Functional connectome fingerprinting: identifying individuals using patterns of brain connectivity. Nat. Neurosci. 18, 1664–1671 (2015).

15. T. Kaufmann, D. Alnæs, N. T. Doan, C. L. Brandt, O. A. Andreassen, L. T. Westlye, Delayed stabilization and individualization in connectome development are related to psychiatric disorders. Nat. Neurosci. 20, 513–515 (2017).

16. O. Miranda-Dominguez, B. D. Mills, S. D. Carpenter, K. A. Grant, C. D. Kroenke, J. T. Nigg, D. A. Fair, Connectotyping: model based fingerprinting of the functional connectome. PloS One. 9, e111048 (2014).

17. M. Fraschini, A. Hillebrand, M. Demuru, L. Didaci, G. L. Marcialis, An EEG-Based Biometric System Using Eigenvector Centrality in Resting State Brain Networks. IEEE Signal Process. Lett. 22, 666–670 (2015).

18. W. Kong, L. Wang, S. Xu, F. Babiloni, H. Chen, EEG Fingerprints: Phase Synchronization of EEG Signals as Biomarker for Subject Identification. IEEE Access. 7, 121165–121173 (2019).

19. D. L. Rocca, P. Campisi, B. Vegso, P. Cserti, G. Kozmann, F. Babiloni, F. D. V. Fallani, Human Brain Distinctiveness Based on EEG Spectral Coherence Connectivity. IEEE Trans. Biomed. Eng. 61, 2406– 2412 (2014).

20. J. de Souza Rodrigues, F. L. Ribeiro, J. R. Sato, R. C. Mesquita, C. E. B. Júnior, Identifying individuals using fNIRS-based cortical connectomes. Biomed. Opt. Express. 10, 2889–2897 (2019).

21. S. Greene, S. Gao, D. Scheinost, R. T. Constable, Task-induced brain state manipulation improves prediction of individual traits. Nat. Commun. 9, 2807 (2018).

22. M. D. Rosenberg, D. Scheinost, A. S. Greene, E. W. Avery, Y. H. Kwon, E. S. Finn, R. Ramani, M. Qiu, R. T. Constable, M. M. Chun, Functional connectivity predicts changes in attention observed across minutes, days, and months. Proc. Natl. Acad. Sci. U. S. A. 117, 3797–3807 (2020).

23. M. Yamashita, Y. Yoshihara, R. Hashimoto, N. Yahata, N. Ichikawa, Y. Sakai, T. Yamada, N. Matsukawa, G. Okada, S. C. Tanaka, K. Kasai, N. Kato, Y. Okamoto, B. Seymour, H. Takahashi, M. Kawato, H. Imamizu, A prediction model of working memory across health and psychiatric disease using whole-brain functional connectivity. eLife. 7 (2018), doi:10.7554/eLife.38844.

24. K. Yoo, M. D. Rosenberg, W.-T. Hsu, S. Zhang, C.-S. R. Li, D. Scheinost, R. T. Constable, M. M. Chun, Connectome-based predictive modeling of attention: Comparing different functional connectivity features and prediction methods across datasets. NeuroImage. 167, 11–22 (2018).

25. E. Bullmore, O. Sporns, The economy of brain network organization. Nat. Rev. Neurosci. 13, 336–349 (2012).

26. S. M. Smith, D. Vidaurre, C. F. Beckmann, M. F. Glasser, M. Jenkinson, K. L. Miller, T. E. Nichols, E. C. Robinson, G. Salimi-Khorshidi, M. W. Woolrich, D. M. Barch, K. Uğurbil, D. C. Van Essen, Functional connectomics from resting-state fMRI. Trends Cogn. Sci. 17, 666–682 (2013).

27. S. Baillet, Magnetoencephalography for brain electrophysiology and imaging. Nat. Neurosci. 20, 327–339 (2017).

28. E. Başar, Chaos in Brain Function: Containing Original Chapters by E. Basar and T. H. Bullock and Topical Articles Reprinted from the Springer Series in Brain Dynamics (Springer Science & Business Media, 1990).

29. R. B. Stein, E. R. Gossen, K. E. Jones, Neuronal variability: noise or part of the signal? Nat. Rev. Neurosci. 6, 389–397 (2005).

30. L. Q. Uddin, Bring the Noise: Reconceptualizing Spontaneous Neural Activity. Trends Cogn. Sci. 24, 734–746 (2020).

31. E. Florin, S. Baillet, The brain’s resting-state activity is shaped by synchronized cross-frequency coupling of neural oscillations. NeuroImage. 111, 26–35 (2015).

32. L. Iemi, N. A. Busch, A. Laudini, S. Haegens, J. Samaha, A. Villringer, V. V. Nikulin, Multiple mechanisms link prestimulus neural oscillations to sensory responses. eLife. 8 (2019), doi:10.7554/eLife.43620.

33. J. Samaha, L. Iemi, S. Haegens, N. A. Busch, Spontaneous Brain Oscillations and Perceptual Decision-Making. Trends Cogn. Sci. 24, 639–653 (2020).

34. S. Bodenmann, T. Rusterholz, R. Dürr, C. Stoll, V. Bachmann, E. Geissler, K. Jaggi-Schwarz, H.-P. Landolt, The functional Val158Met polymorphism of COMT predicts interindividual differences in brain alpha oscillations in young men. J. Neurosci. Off. J. Soc. Neurosci. 29, 10855–10862 (2009).

35. S. Haegens, H. Cousijn, G. Wallis, P. J. Harrison, A. C. Nobre, Inter-and intra-individual variability in alpha peak frequency. NeuroImage. 92, 46–55 (2014).

36. S. Baillet, J. C. Mosher, R. M. Leahy, Electromagnetic brain mapping. IEEE Signal Process. Mag. 18, 14–30 (2001).

37. R. S. Desikan, F. Ségonne, B. Fischl, B. T. Quinn, B. C. Dickerson, D. Blacker, R. L. Buckner, A. M. Dale, R. P. Maguire, B. T. Hyman, M. S. Albert, R. J. Killiany, An automated labeling system for subdividing the human cerebral cortex on MRI scans into gyral based regions of interest. NeuroImage. 31, 968– 980 (2006).

38. P. E. Shrout, J. L. Fleiss, Intraclass correlations: Uses in assessing rater reliability. Psychol. Bull. 86, 420–428 (1979).

39. R. McIntosh, B. Mišić, Multivariate Statistical Analyses for Neuroimaging Data. Annu. Rev. Psychol. 64, 499–525 (2013).

40. T. Yeo, Thomas, F. M. Krienen, J. Sepulcre, M. R. Sabuncu, D. Lashkari, M. Hollinshead, J. L. Roffman, J. W. Smoller, L. Zöllei, J. R. Polimeni, B. Fischl, H. Liu, R. L. Buckner, The organization of the human cerebral cortex estimated by intrinsic functional connectivity. J. Neurophysiol. 106, 1125– 1165 (2011).

41. S. Noble, M. N. Spann, F. Tokoglu, X. Shen, R. T. Constable, D. Scheinost, Influences on the Test-Retest Reliability of Functional Connectivity MRI and its Relationship with Behavioral Utility. Cereb. Cortex N. Y. N 1991. 27, 5415–5429 (2017).

42. J. Cabral, M. L. Kringelbach, G. Deco, Functional connectivity dynamically evolves on multiple time-scales over a static structural connectome: Models and mechanisms. NeuroImage. 160, 84–96 (2017).

43. M. Nentwich, L. Ai, J. Madsen, Q. K. Telesford, S. Haufe, M. P. Milham, L. C. Parra, Functional connectivity of EEG is subject-specific, associated with phenotype, and different from fMRI. NeuroImage. 218, 117001 (2020).

44. C. L. Horien, X. Shen, D. Scheinost, R. T. Constable, The individual functional connectome is unique and stable over months to years. Neuroimage (2019), doi:10.1016/j.neuroimage.2019.02.002.

45. M. J. Brookes, M. Woolrich, H. Luckhoo, D. Price, J. R. Hale, M. C. Stephenson, G. R. Barnes, S. M. Smith, P. G. Morris, Investigating the electrophysiological basis of resting state networks using magnetoencephalography. Proc. Natl. Acad. Sci. U. S. A. 108, 16783–16788 (2011).

46. B. A. E. Hunt, P. K. Tewarie, O. E. Mougin, N. Geades, D. K. Jones, K. D. Singh, P. G. Morris, P. A. Gowland, M. J. Brookes, Relationships between cortical myeloarchitecture and electrophysiological networks. Proc. Natl. Acad. Sci. 113, 13510 (2016).

47. G. Michalareas, J. Vezoli, S. van Pelt, J.-M. Schoffelen, H. Kennedy, P. Fries, Alpha-Beta and Gamma Rhythms Subserve Feedback and Feedforward Influences among Human Visual Cortical Areas. Neuron. 89, 384–397 (2016).

48. B. Morillon, S. Baillet, Motor origin of temporal predictions in auditory attention. Proc. Natl. Acad. Sci. 114, E8913–E8921 (2017).

49. S. Haufe, P. DeGuzman, S. Henin, M. Arcaro, C. J. Honey, U. Hasson, L. C. Parra, Elucidating relations between fMRI, ECoG, and EEG through a common natural stimulus. NeuroImage. 179, 79–91 (2018).

50. N. K. Logothetis, J. Pauls, M. Augath, T. Trinath, A. Oeltermann, Neurophysiological investigation of the basis of the fMRI signal. Nature. 412, 150–157 (2001).

51. J. F. Nottage, J. Horder, State-of-the-Art Analysis of High-Frequency (Gamma Range) Electroencephalography in Humans. Neuropsychobiology. 72, 219–228 (2015).

52. E. M. Whitham, K. J. Pope, S. P. Fitzgibbon, T. Lewis, C. R. Clark, S. Loveless, M. Broberg, A. Wallace, D. DeLosAngeles, P. Lillie, A. Hardy, R. Fronsko, A. Pulbrook, J. O. Willoughby, Scalp electrical recording during paralysis: Quantitative evidence that EEG frequencies above 20Hz are contaminated by EMG. Clin. Neurophysiol. 118, 1877–1888 (2007).

53. S. Yuval-Greenberg, O. Tomer, A. S. Keren, I. Nelken, L. Y. Deouell, Transient induced gamma-band response in EEG as a manifestation of miniature saccades. Neuron. 58, 429–441 (2008).

54. Y. Bagherzadeh, D. Baldauf, D. Pantazis, R. Desimone, Alpha Synchrony and the Neurofeedback Control of Spatial Attention. Neuron. 105, 577-587.e5 (2020).

55. M. S. Clayton, N. Yeung, R. Cohen Kadosh, The many characters of visual alpha oscillations. Eur. J. Neurosci. 48, 2498–2508 (2018).

56. J. J. Foster, E. Awh, The role of alpha oscillations in spatial attention: limited evidence for a suppression account. Curr. Opin. Psychol. 29, 34–40 (2019).

57. T. Lennert, S. Samiee, S. Baillet, Coupled oscillations enable rapid temporal recalibration to audiovisual asynchrony. Commun. Biol. 4, 559 (2021).

58. J. C. Mosher, S. Baillet, R. M. Leahy, in IEEE Workshop on Statistical Signal Processing, 2003 (2003), pp. 294–297.

59. E. Sareen, S. Zahar, D. Van De Ville, A. Gupta, A. Griffa, E. Amico, “Exploring MEG brain fingerprints: evaluation, pitfalls, and interpretations” (preprint, Neuroscience, 2021), doi:10.1101/2021.02.15.431253.

60. S. Sadaghiani, M. J. Brookes, S. Baillet, “Connectomics of Human Electrophysiology” (preprint, PsyArXiv, 2021), doi:10.31234/osf.io/dr7zh.

61. M. D. Rosenberg, E. S. Finn, D. Scheinost, X. Papademetris, X. Shen, R. T. Constable, M. M. Chun, A neuromarker of sustained attention from whole-brain functional connectivity. Nat. Neurosci. 19, 165–171 (2016).

62. M. D. Rosenberg, E. S. Finn, D. Scheinost, R. T. Constable, M. M. Chun, Characterizing Attention with Predictive Network Models. Trends Cogn. Sci. 21, 290–302 (2017).

63. T. Harmelech, R. Malach, Neurocognitive biases and the patterns of spontaneous correlations in the human cortex. Trends Cogn. Sci. 17, 606–615 (2013).

64. H. Cai, J. Zhu, Y. Yu, Robust prediction of individual personality from brain functional connectome. Soc. Cogn. Affect. Neurosci. 15, 359–369 (2020).

65. C. Parkinson, A. M. Kleinbaum, T. Wheatley, Similar neural responses predict friendship. Nat. Commun. 9, 332 (2018).

66. D. C. Glahn, A. M. Winkler, P. Kochunov, L. Almasy, R. Duggirala, M. A. Carless, J. C. Curran, R. L. Olvera, A. R. Laird, S. M. Smith, C. F. Beckmann, P. T. Fox, J. Blangero, M. E. Raichle, Genetic Control over the Resting Brain. Proc. Natl. Acad. Sci. U. S. A. 107, 1223–1228 (2010).

67. M. S. Korgaonkar, K. Ram, L. M. Williams, J. M. Gatt, S. M. Grieve, Establishing the resting state default mode network derived from functional magnetic resonance imaging tasks as an endophenotype: A twins study. Hum. Brain Mapp. 35, 3893–3902 (2014).

68. O. Miranda-Dominguez, E. Feczko, D. S. Grayson, H. Walum, J. T. Nigg, D. A. Fair, Heritability of the human connectome: A connectotyping study. Netw. Neurosci. Camb. Mass. 2, 175–199 (2018).

69. C. A. Hodgkinson, M.-A. Enoch, V. Srivastava, J. S. Cummins-Oman, C. Ferrier, P. Iarikova, S. Sankararaman, G. Yamini, Q. Yuan, Z. Zhou, B. Albaugh, K. V. White, P.-H. Shen, D. Goldman, Genome-wide association identifies candidate genes that influence the human electroencephalogram. Proc. Natl. Acad. Sci. U. S. A. 107, 8695–8700 (2010).

70. E. Leppäaho, H. Renvall, E. Salmela, J. Kere, R. Salmelin, S. Kaski, Discovering heritable modes of MEG spectral power. Hum. Brain Mapp. 40, 1391–1402 (2019).

71. E. Salmela, H. Renvall, J. Kujala, O. Hakosalo, M. Illman, M. Vihla, E. Leinonen, R. Salmelin, J. Kere, Evidence for genetic regulation of the human parieto-occipital 10-Hz rhythmic activity. Eur. J. Neurosci. 44, 1963–1971 (2016).

72. T. Kaufmann, D. Alnæs, C. L. Brandt, F. Bettella, S. Djurovic, O. A. Andreassen, L. T. Westlye, Stability of the Brain Functional Connectome Fingerprint in Individuals With Schizophrenia. JAMA Psychiatry. 75, 749 (2018).

73. F. Tadel, S. Baillet, J. C. Mosher, D. Pantazis, R. M. Leahy, Brainstorm: a user-friendly application for MEG/EEG analysis. Comput. Intell. Neurosci. 2011, 879716 (2011).

74. J. Gross, S. Baillet, G. R. Barnes, R. N. Henson, A. Hillebrand, O. Jensen, K. Jerbi, V. Litvak, B. Maess, R. Oostenveld, L. Parkkonen, J. R. Taylor, V. van Wassenhove, M. Wibral, J.-M. Schoffelen, Good practice for conducting and reporting MEG research. NeuroImage. 65, 349–363 (2013).

75. B. Fischl, FreeSurfer. NeuroImage. 62, 774–781 (2012).

76. Bruns, R. Eckhorn, H. Jokeit, A. Ebner, Amplitude envelope correlation detects coupling among incoherent brain signals. Neuroreport. 11, 1509–1514 (2000).

77. P. Welch, The use of fast Fourier transform for the estimation of power spectra: A method based on time averaging over short, modified periodograms. IEEE Trans. Audio Electroacoustics. 15, 70–73 (1967).

78. R Core Team, R: A Language and Environment for Statistical Computing (R Foundation for Statistical Computing, Vienna, Austria, 2020; https://www.R-project.org/).

79. M. Mowinckel, D. Vidal-Piñeiro, ggseg: Plotting Tool for Brain Atlases (2021; https://CRAN.R-project.org/package=ggseg).

80. R. McIntosh, N. J. Lobaugh, Partial least squares analysis of neuroimaging data: applications and advances. NeuroImage. 23 Suppl 1, S250–263 (2004).

## References

1. E. Amico, J. Goñi, The quest for identifiability in human functional connectomes. Sci. Rep. 8, 8254 (2018).

2. E. S. Finn, X. Shen, D. Scheinost, M. D. Rosenberg, J. Huang, M. M. Chun, X. Papademetris, R. T. Constable, Functional connectome fingerprinting: identifying individuals using patterns of brain connectivity. Nat. Neurosci. 18, 1664–1671 (2015).

3. T. Yeo, Thomas, F. M. Krienen, J. Sepulcre, M. R. Sabuncu, D. Lashkari, M. Hollinshead, J. L. Roffman, J. W. Smoller, L. Zöllei, J. R. Polimeni, B. Fischl, H. Liu, R. L. Buckner, The organization of the human cerebral cortex estimated by intrinsic functional connectivity. J. Neurophysiol. 106, 1125–1165 (2011).

4. R. S. Desikan, F. Ségonne, B. Fischl, B. T. Quinn, B. C. Dickerson, D. Blacker, R. L. Buckner, A. M. Dale, R. P. Maguire, B. T. Hyman, M. S. Albert, R. J. Killiany, An automated labeling system for subdividing the human cerebral cortex on MRI scans into gyral based regions of interest. NeuroImage. 31, 968– 980 (2006).

